# H_2_O_2_-induced astrocytic collagen triggers neuronal death via fucosylation-dependent glial barrier formation upon ischemic stroke

**DOI:** 10.1101/2025.05.01.651594

**Authors:** Jae-Hun Lee, Hyun Jun Jang, In-Young Hwang, Jiwoon Lim, Jiwoong Jung, Myungju Kim, Eunji Lee, Woojin Won, Elijah Hwejin Lee, Ki Jung Kim, Ki Duk Park, Woonggyu Jung, Boyoung Lee, Seungjun Ryu, C. Justin Lee

## Abstract

The cascade of molecular and cellular events leading to neuronal death after a focal ischemic stroke remains enigmatic. Although astrocytes form glial barriers that may protect surrounding tissue, how barriers develop and contribute to neuronal death is unclear. Here, we show that H₂O₂ induces astrocytic type I collagen (COL1)-production via miR-29-mediated post-transcriptional and fucosylation-dependent post-translational regulations, leading to integrin activation and neuronal death. In photothrombosis (PT)-induced cortical stroke model, PT triggered H₂O₂-surge, astrogliosis, glial barrier formation, COL1 expression, fibrotic scarring, altered N-glycosylation, neuronal loss, and neurological deficits. Remarkably, these effects were reversed by astrocyte-specific COL1 or FUT8 gene-silencing or treatment with KDS12025, an H₂O₂-decomposing peroxidase enhancer. KDS12025’s neuroprotective effects were also recapitulated in non-human primate stroke model. These findings delineate a previously unrecognized astrocyte-driven mechanism in which oxidative stress induces COL1 production, promoting neuronal death, and position H₂O₂ and astrocytic COL1 and FUT8 as promising therapeutic targets for ischemic stroke.

## Introduction

Reactive oxygen species (ROS) are essential for brain physiology and function. They play a dual role, acting as critical signaling molecules at physiological levels while contributing to oxidative damage when excessively produced. Under normal conditions, ROS regulate synaptic plasticity^1^, neuronal differentiation^2^, and immune responses^3^, contributing to essential brain processes such as learning and memory^4^. However, when ROS levels exceed the capacity of antioxidant defense systems, they can induce oxidative stress, leading to lipid peroxidation^5^, protein oxidation^6^, and DNA damage^7^. The brain is particularly vulnerable to oxidative stress due to its abundant polyunsaturated fatty acids^8^, and relatively low antioxidant capacity^9^. As a result, ROS have been implicated in the initiation and progression of various neurodegenerative diseases, including ischemic stroke^10^, Alzheimer’s disease^11^, and Parkinson’s disease^12^. In these conditions, excessive ROS generation can trigger neuroinflammation, blood-brain barrier disruption, and neuronal apoptosis, exacerbating disease pathology. However, how specific brain cells respond to ROS and how this affects neuronal survival remain unclear.

Among the various brain cell types, astrocytes are pivotal in maintaining brain homeostasis and undergo structural and functional alterations in response to brain injuries to transform into reactive astrocytes^13,14^. Especially after an ischemic stroke, these reactive astrocytes tightly line up and surround the injured area to form a glial barrier, presumably to protect the uninjured area from further damage^15^. These reactive astrocytes contribute to extracellular matrix (ECM) remodeling by producing and secreting glycoproteins associated with ECM and collagen fibril organization^16,17^, which can be dynamically regulated in response to injury^18,19^. Notably, our recent studies highlight that excessive H_2_O_2_ produced by monoamine oxidase B (MAO-B) in severe reactive astrocytes plays a crucial role in neurodegeneration in Alzheimer’s disease^11^. However, the molecular mechanisms underlying the interactions between astrocytic H_2_O_2_ and ECM and between ECM and neurons and their impact on neurodegeneration are entirely unknown.

In this study, we set out to test the idea that severe reactive astrocytes are involved in neuronal death after ischemic stroke. Through transcriptomic analysis using RNA and ATAC-seq, we found that H₂O₂ induces the upregulation of genes related to ECM production, including type I collagen (COL1) from astrocytes, alongside genes regulating N-linked glycosylation, a key post-translational modification essential for ECM assembly. We further identified that the production of COL1 is modulated by post-transcriptional regulation and that miR-29 is involved in this process. Utilizing the photothrombosis (PT) -induced acute ischemic stroke models of mouse and non-human primate (NHP), we demonstrate that PT-induced H₂O₂ and H₂O₂-induced severe reactive astrocytes produce toxic collagen through N-linked glycosylation, significantly contributing to ECM remodeling, glial barrier formation, neuronal death, and motor impairment. Treatment with the H₂O₂-decomposing peroxidase enhancer KDS12025 and astrocytic COL1 or FUT8 gene-silencing successfully reverses the pathological cascade of stroke and motor impairment, highlighting a key aspect of astrocytic COL1 in neuronal death. To our knowledge, this is the first study to comprehensively dissect the astrocyte-mediated pathological cascade underlying stroke and neurological deficits.

## Results

### H_2_O_2_ turns on collagen production in cortical astrocytes via post-transcriptional and post-translational regulations

Astrocytes, the primary glial cells responsible for maintaining homeostasis, become reactive in response to oxidative stress, undergoing functional changes that influence ECM remodeling and scar formation^20,21^. However, the molecular mechanisms by which H₂O₂ modulates astrocyte-derived ECM components remain incompletely understood. To investigate the direct involvement of H2O2 in astrocyte-derived ECM remodeling, we performed RNA-seq to analyze the mRNA expression profiles after 3 hours of 200 µM H2O2 treatment on cortical astrocytes (Figure 1A). Among the significantly changed genes (p-values < 0.05, log2(FC) > 0.6), 11.5% were identified, exhibiting upregulation (42.5%) or downregulation (57.5%) one day after H2O2 treatment (D1). Notably, only 1.15% of the total genes demonstrated significant expression changes three days after H2O2 treatment (D3) (Figures 1B, S1A, and S1B), suggesting that the gene expression changes induced by 3 hours of H2O2 exposure are reversible by D3. Hence, we used differentially expressed genes (DEGs) of D1 for further gene ontology (GO) and pathway analysis, utilizing Reactome and KEGG databases. Terms associated with collagen metabolism and collagen-containing extracellular matrix were highly enriched, particularly among up-regulated genes: Gene set enrichment analysis (GSEA) also showed a positive correlation in collagen metabolism and collagen-containing ECM (Figures 1C, 1D, and S1C-G). Indeed, expression of *Col1α1* and *Col1α2*, which encode the alpha-1 and -2 subunits of COL1, increased significantly in D1 after H_2_O_2_ treatment and reduced in D3 (Figure 1E). However, at the protein level, we observed a gradual increase of COL1, peaking at 72 hours (D3) after H_2_O_2_ treatment and returning to control levels at 168 hours (D7) (Figures 1F and G). This discrepancy in the time courses of mRNA and protein expression implies that there might be post-translational modifications (PTM) affecting protein stability or function of COL1. In support, the GO analysis shows the enrichment of PTM-related genes, including N-linked glycosylation and positive regulation of PTM genes (Figures 1H and S1H). These results raise the possibility that PTM of COL1 and related proteins, particularly N-linked glycosylations, might be essential in regulating collagen stability during ECM remodeling in barrier-forming reactive astrocytes. Together, these findings imply that astrocytes turn on collagen production upon H_2_O_2_-treatment in two independent ways: 1) by increasing mRNA expression of collagen-related genes and 2) by PTM of COL1-related proteins, possibly via N-linked glycosylation.

**Figure 1.**
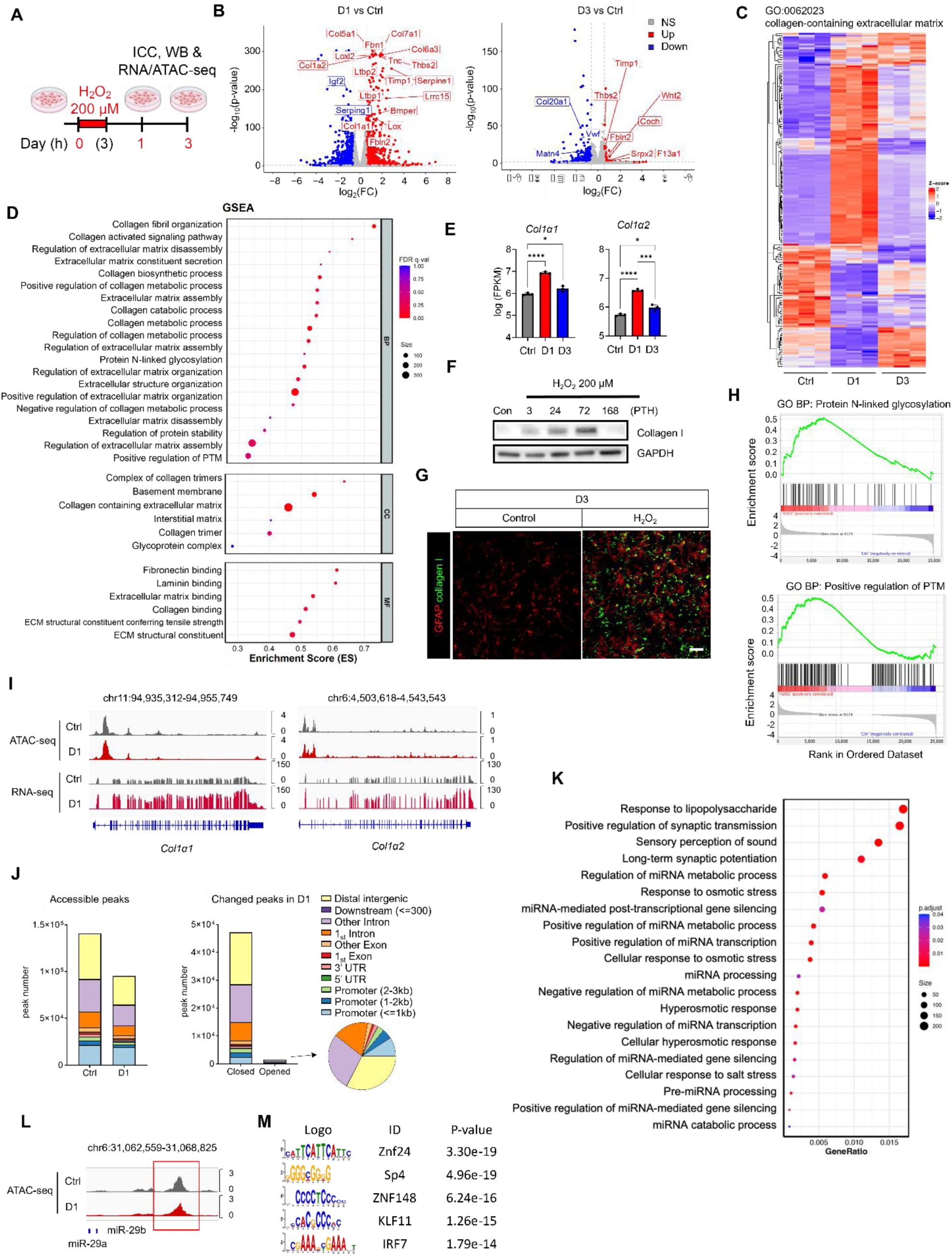
H_2_O_2_ induces collagen production via post-transcriptional regulation by miR-29 and post-translational regulation by N-linked glycosylation in cortical astrocytes. (A) Schematic image of experimental schedule (B) Volcano plot of RNA-seq experiment, genes with p<0.05, log2(FC)>|0.6|) in GO:0062023, collagen containing extracellular matrix are indicated, Red: Up-regulated genes, Blue: Down-regulated genes (C) Heat map of gene expressions related to GO terms associated with collagen-containing extracellular matrix (D) GO analysis of RNA-seq data from H_2_O_2_ treated cortical astrocyte compared with control, BP: Biological process CC: Cellular component MF: Molecular function (E) Log (FPKM) of Col1α1 and *Col1α2* genes from RNA-seq data (F) Western blot result of collagen I protein expression in H_2_O_2_ treated cortical astrocyte (PTH: post treatment hour) (G) Representative ICC image with GFAP and collagen I in cultured astrocyte at D3 (H) GO enrichment analysis of genes in GO:0006487, protein N-linked glycosylation and GO:0043687, post-translational modifications (I) Number and proportion of accessible peaks and differentially regulated peaks of *Col1α1* and *Col1α2* (J) Integrative genome viewer (IGV) images of Col1α1 and Col1α2 from RNA- and ATAC-seq (x-axis; the genome track of indicated genes, y-axis; read count per million (CPM)) (K) GO: BP of Closed peaks in D1 (L) Up: IGV images of miR-29a and miR-29b regulatory regions from ATAC-seq (x-axis; the genome track of miR-29s, y-axis; CPM) (M) Motif enrichment analysis of the regulatory regions of miR-29a and miR-29b (regions in (L) red box) using JASPAR (2024)

To delineate the epigenetic mechanisms of how H_2_O_2_ increases mRNA expression of collagen-related genes in astrocytes, we performed the assay of transposase-accessible chromatin sequencing (ATAC-seq). Contrary to expectations, the total number of accessible peaks was reduced to 68% compared to the control (from 140,713 to 95,460 peaks); at the same time, accessible peaks in promoters changed minimally (from 30,351 to 24,805 peaks) (Figure 1I). Unlike the increased mRNA levels from RNA-seq results, accessible peaks in *Col1α1* and *Col1α2* promoter regions from ATAC-seq results showed no significant difference between with and without H_2_O_2_ treatment (Figure 1I), strongly suggesting a post-transcriptional regulation. Indeed, accessibility was markedly closed in distal intergenic regions and introns, where clusters of non-coding genes such as microRNAs (miRNAs)^22^ reside (Figure 1J), suggesting that H_2_O_2_ treatment triggers the down-regulation of these non-coding miRNAs. We performed GO analysis on the closed peaks after H_2_O_2_ treatment (D1) and found an enrichment of miRNA processing genes (Figure 1K). This indicates that chromatin regions involved in miRNA processes are closed after H_2_O_2_ treatment. Because miR-29a has been reported to target and inhibit collagen expression possibly by degrading *Col1α1* mRNA^23^, we conducted an in-depth analysis and identified the miR-29a/b/c target sites in 3’UTR of *Col1α1* regions using TargetScan^24^ (Figure S1I). More importantly, accessible peaks of miR-29a and miR-29b showed a marked closure of the regulatory regions after H_2_O_2_ treatment D1 (Figure 1L), raising a possible disinhibitory role of miR-29a and miR-29b on *Col1α1* mRNA expression. Further analysis of the differentially accessible peaks to investigate transcription factor binding motifs and signaling pathways revealed the multiple motifs that are prevalent in the intergenic regulatory regions of miR-29a and miR-29b (red box in Figure 1L), including transcription factors linked to ROS, inflammation, and chromatin remodelers (Figure 1K). Collectively, these results suggest that H_2_O_2_ triggers collagen production in cortical astrocytes via disinhibitory post-transcriptional regulation of *Col1α1* and *Col1α2* mRNAs by closing regulatory regions of miR-29a/b/c, providing an alternative therapeutic target to inhibit astrocytic collagen production.

### H_2_O_2_-induced astrocytic Col1 induces neuronal death through integrin receptor activation

To investigate the possible role of H_2_O_2_-induced astrocytic COL1 on cortical neurons, we performed a sandwich culture (Figure 2A-D). Primary cortical neurons and astrocytes were seeded separately on a culture dish and an insert. After inducing the collagen production of astrocytes by H_2_O_2_ treatment, the insert was placed on top of cortical neurons grown in a culture dish (Figure 2A). After 2 days of sandwich culture, we found a significant reduction in the number of live cortical neurons cultured with H_2_O_2_-treated astrocytes compared to those with untreated astrocytes (Figure 2B), suggesting a release of neurotoxic COL1 from the H_2_O_2_-treated astrocytes. To test if astrocytic COL1 is necessary for H_2_O_2_-induced neuronal death, we developed *Col1α1*-shRNA to gene-silence *Col1α1* in cultured astrocytes (Figures S2A-C). A sandwich culture of cortical neurons and *Col1α1* gene-silenced astrocytes treated with H_2_O_2_ showed significantly reduced neuronal death compared to the control Scrambled (Sc)-shRNA group (Figure 2C and D), indicating that astrocytic COL1 is necessary for H_2_O_2_-induced neurotoxicity. Indeed, direct application of 1 µg/ml COL1 to cultured neurons killed neurons significantly within one day and almost completely after 3-day treatment (Figure 3E and F). These results indicate that H_2_O_2_-induced astrocytic COL1 triggers slow neuronal death, which takes up to three days.

**Figure 2.**
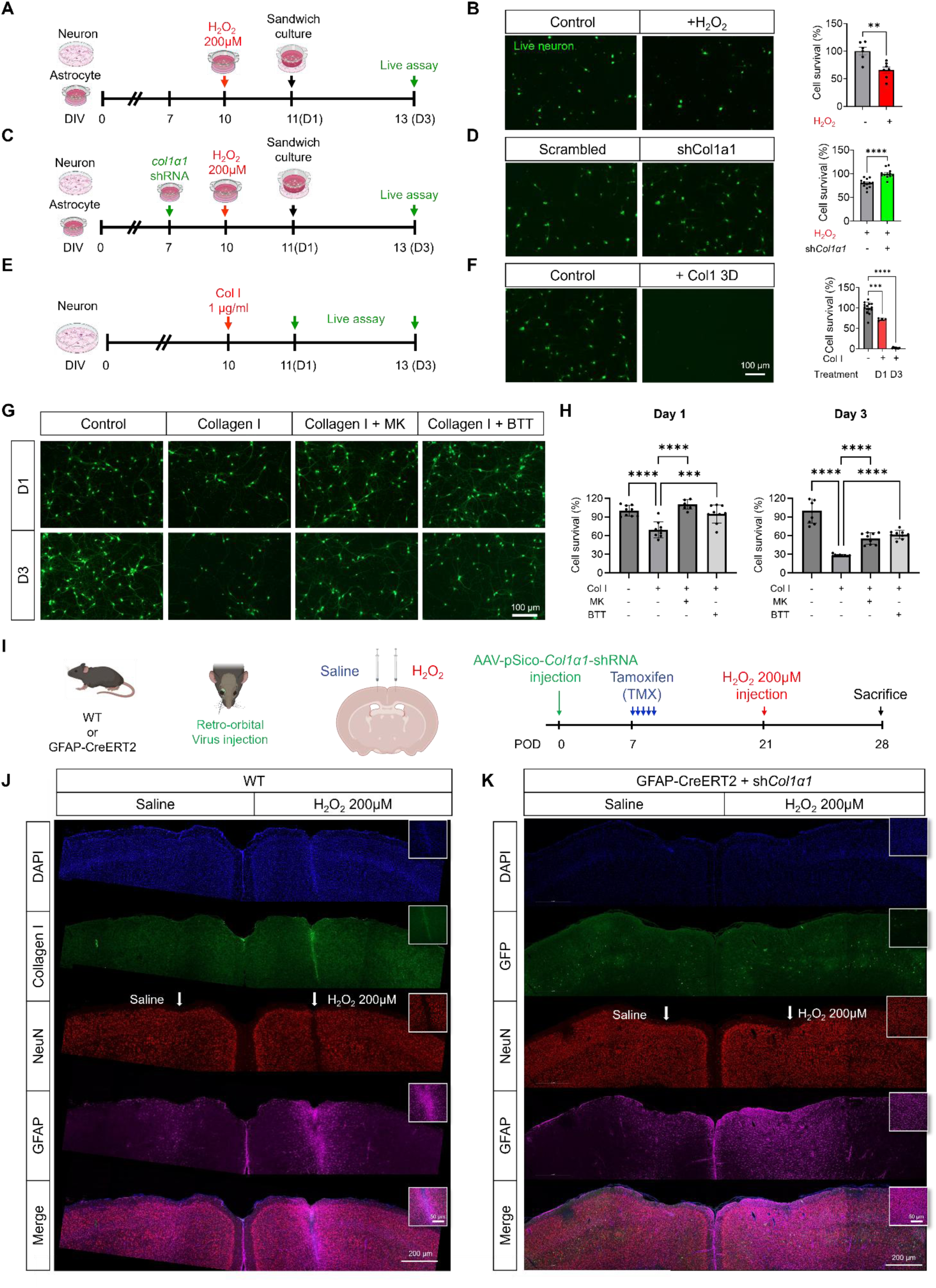
H_2_O_2_-induced astrocytic COL1 triggers neuronal death through integrin receptor activation. (A to F) Experimental schedule for cortical neuron and astrocyte sandwich culture with ICC image and quantification (error bar: SEM, Mann-Whitney U test, \*\**p*<0.01) (E and F) Experimental schedule for collagen I treatment on cortical neuron and ICC image with live neuron quantification (error bar: SEM, Paired, One-way ANOVA with Dunnett’s post hoc, \*\*\**p*<0.001, \*\*\*\**p*<0.0001) (G) Representative image of neurons treated with collagen I and integrin inhibitors (H) Quantification of cell survival treated with type I collagen and integrin inhibitors (One-way ANOVA with Dunnett’s post hoc, \*\*\**p*<0.001, \*\*\*\**p*<0.0001) (I) Experimental schedule and Injection route of virus (J) Represent image of DAPI (blue), Collagen I (green), NeuN (red), and GFAP (magenta) after saline and H_2_O_2_ injection (inset image indicate injection site) (K) Represent image of DAPI (blue), GFP (green), NeuN (red), and GFAP (magenta) with *col1α1* shRNA after saline and H_2_O_2_ injection

**Figure 3.**
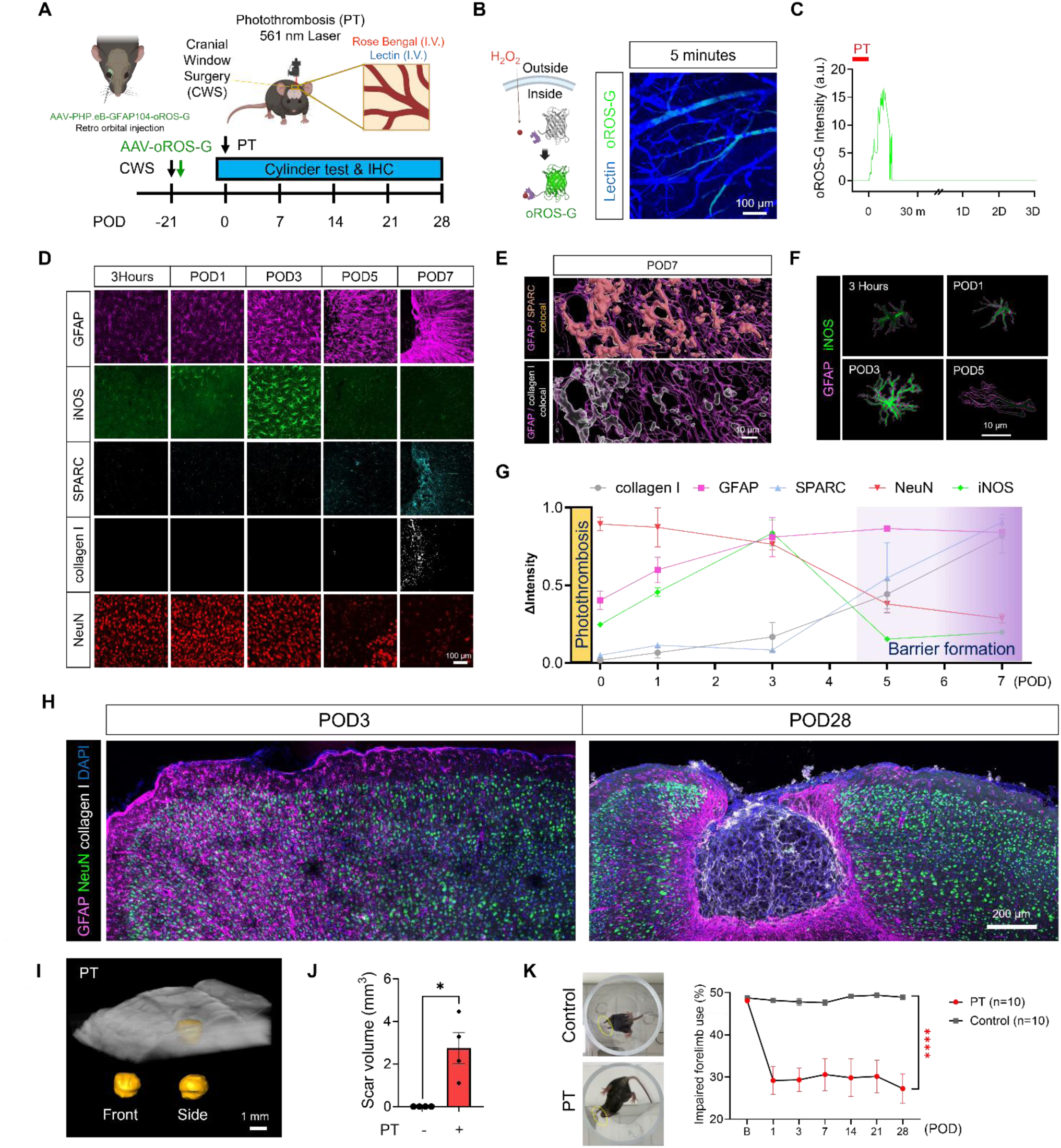
PT induces H_2_O_2_ surge, severe astrogliosis, glial barrier, collagen production, fibrotic scar, neuronal death, and motor impairment. (A) Schematic images of PT modeling and the timeline of experiment (POD: post operation day, Rose Bengal, 15 mg/ml, 20 µl, intra-venous (I.V.), Lectin, 1 mg/ml, 20 µl, I.V., AAV-PHP.eB-GFAP104-oROS-G, 20 µl) (B) Schematic image represent mode of action of the oROS-G and intra-vital imaging of blood vessel (Lectin, blue) and H_2_O_2_ sensor (oROS-G, green) (C) Quantification of oROS signals after PT modeling (Red bar: duration of thrombosis, PT: Photothrombosis, m: minutes, D: day) (D) Time course protein expression changes of GFAP, iNOS, SPARC, collagen I, and NeuN at infarct region (E) Imaris reconstruction for colocalization of GFAP with collagen I and SPARC at POD7 (F) Reconstructed images of GFAP (magenta) with iNOS (green) at 3 hours, POD1, 3, and 5 (G) Quantification of various protein expressions after PT (H) Representative images of GFAP (magenta), collagen I (white), NeuN (green) with DAPI (blue) at infarct region in POD3 and 28 (I) SOCM image and 3D reconstruction of fibrotic scar (J) Quantification of scar volume in PT group (Contra: contralateral side, error bar: SEM, Mann-Whitney’s U test, \**p*<0.05) (K) Representative image of cylinder test with paw touches (yellow dotted circle) and ratio of impaired forelimb use (Two-way ANOVA with *Dunnett’s* post hoc, **** *p*<0.0001, *** *p*<0.001, error bar: SEM, B: baseline, POD: post operation day)

Collagen protein can act as a signaling molecule by interacting with several receptors, including integrins, receptor tyrosine kinases, and discoidin domain receptors^25^. Cortical neurons have various types of integrins that interact with the ECM, and these could be possible receptors of astrocytic COL1, leading to cell death signaling. To test this possibility, we treated cortical neurons with MK-0429, a small molecule integrin inhibitor, or BTT-3033, another integrin inhibitor, 30 minutes before COL1 treatment and counted the number of surviving cells (Figures 2G and H). Quantification for cell survival showed that both integrin inhibitors significantly reduce COL1-induced neuronal death (Figure 2H). These results indicate that neuronal integrins are necessary for COL1-induced neuronal death in the cortical neurons.

To test if H_2_O_2_ is sufficient to induce COL1 and neuronal death *in vivo*, we injected H_2_O_2_ directly into the cortex (Figures 2I). We found that 200 µM H_2_O_2_, but not 10 or 100 µM H_2_O_2_, was sufficient to trigger COL1 production and neuronal death (Figures 2J and S3D), which were all prevented by gene-silencing of astrocytic *Col1α1* (Figures 2K). Together, these results indicate that H_2_O_2_-induced astrocytic COL1 triggers a slow neuronal death through integrins, providing a possible explanation for the slow neuronal loss in neurodegenerative diseases.

### Photothrombosis (PT) induces H_2_O_2_-surge, severe astrogliosis, glial barrier, collagen production, fibrotic scar, neuronal death, and motor impairment

Among various brain diseases, an ischemic stroke, which shows the extensive remodeling of ECM occurs^26^, ROS generation by ischemic stress has been suggested as an initial trigger of pathological cascade^10^. To investigate the involvement of astrocytic ECM modulation in excessive ROS generation condition, we employed a mouse stroke model by inducing PT (Rose Bengal irradiated with 561 nm laser through a cranial window) in the primary motor cortex (M1) and secondary motor cortex (M2) regions of the mouse brain (Figure 3A), as previously described^27^. The blood flow through the vessels was monitored via lectin, allowing visualization of the PT-induced infarction (Video S1). To monitor the cytosolic H_2_O_2_ level in astrocytes, we subjected the mice to retro-orbital injection of the adeno-associated virus (AAV-PHP.eB-GFAP104-oROS-G) carrying the recently developed green H_2_O_2_-specific sensor (oROS-G)^28^ for astrocyte-specific expression (Figures 3B and S4A and B). The virus was packaged in an AAV serotype PHP.eB to retro-orbitally infect the brain without damaging the meninges and causing undesirable astrogliosis due to needle insertion^29^. We observed a surge of oROS-G signal immediately after PT lasting about 5∼10 minutes in astrocytic endfeet along the blood vessels near the infarct region (Figures 3A, 3C, S4C, S4D, and Video S1). The oROS-G signal disappeared 15 minutes after PT, likely due to PT-induced blood vessel disintegration and astrocytic endfeet detachment (Figure S4D), raising the possibility that the elevated H_2_O_2_level could persist even after 15 minutes. Time-lapse intravital imaging of the infarct region showed permanent disruption of vascular structure at POD1 (Figure S4D) and massive angiogenesis until POD28 (Figures S4G). These results indicate an excessive increase in cytosolic H_2_O_2_levels in astrocytes at the initial stages, leading to permanent vascular disruption and angiogenesis at the infarct region in this model. To investigate the dynamic changes of astrocytes in the infarct region, we performed time-lapse imaging of astrocytes using Aldh1l1-eGFP mice and found that PT causes reduction of eGFP intensity together with vascular fragmentation at the infarct region (Figures S5A and B). Since eGFP protein is sensitive to pH and can be quenched in an acidic environment^30^, this result indicates that excessive H_2_O_2_production in this region could induce metabolic changes in astrocytes, leading to a decrease in pH through local acidification^31^.

To elucidate the time course of molecular and cellular changes in the ECM of the infarct region, we performed immunohistochemistry for various cellular and collagen-related markers at different time points after PT (Figures 3D, S6A, and S6B). As early as 3 hours post-infarction, we observed highly reactivated astrocytes with increased GFAP signals in the infarct region (Figure 3D). Along with astrocytic hypertrophy, expression of iNOS, a marker of severe reactive astrocytes^11^, gradually increased until post-operation day 3 (POD3), then suddenly decreased by POD5 (Figure 3D and F) as astrocytes began to show polarity with extended morphology (Figures S7A-E). Notably, starting from POD5, we observed an appearance of COL1 and secreted protein acidic and rich in cysteine (SPARC), a matricellular collagen-binding protein known to be involved in collagen assembly, secretion^32^, and fibrillogenesis^33^, in barrier-forming reactive astrocytes, coinciding with the disappearance of a neuronal marker NeuN and astrocyte marker GFAP in infarct region (Figures 3D and 3G). Colocalization of COL1 and SPARC with GFAP in the glial barrier was identified with 3D reconstruction analysis (Figure 3E). Surprisingly, most NeuN-positive neurons were still present at POD3 (Figures 3D and 3G), suggesting a possibility that these neurons are still alive. However, by the time of POD28, NeuN-positive neurons were utterly absent in the infarct region and within the glial barrier where COL1 is abundantly expressed (Figures 3H (merged images) and S8F (separated images)). Unlike GFAP, the expression of Iba1, a microglia marker, gradually increased, accumulating only in the infarct region but not in the glial barrier (Figures S6A and B). Consistent with the immunohistochemical evidence, under the intra-vital imaging of GFAPCreERT2:tdTomato mice, we observed an appearance and accumulation of reactive astrocytes near the infarct region from POD7 to POD28, with newly produced collagen fibers as visualized by the second harmonic generation (SHG) Figures S4E-G). These results indicate that ischemic stroke causes an initial surge of H_2_O_2_, which precedes dramatic changes in various protein expressions, leading to reactive astrogliosis, glia barrier formation, collagen production, and neurodegeneration.

To examine the reactivity of astrocytes in terms of structural features within the glial barrier, we conducted detailed morphological analyses in terms of temporal and spatial patterns. Temporally and spatially, GFAP-positive astrocytes displayed diverse morphological features at POD7 and POD28, as well as within the proximal, distal, and outside regions of the glial barrier, with more severe astrogliosis closer to the barrier (Figures S7G-N). Based on these unique morphological features of barrier-forming reactive astrocytes, we performed serial optical coherence microscopy (SOCM) imaging from serial brain sections of the entire brain, as previously described^34^. SOCM allowed convenient 3D reconstruction and delineation of the borders of the fibrotic scar for accurate measurement of the scar volume (Figure 3I). We observed the PT-induced scar formation with an average volume of around 3 mm^3^ at POD28 (Figure 3J). Finally, we monitored the motor function of each mouse at various time points using a cylinder test generally used for monitoring behavioral changes after stroke^35–37^ by calculating the percentage of impaired forelimb use. PT caused an immediate and significant drop in impaired forelimb use, which persisted throughout POD28 (Figure 3K). Taken together, these results indicate that ischemic stroke induced by PT initiates an H_2_O_2_-surge in astrocytes, followed by a cascade of pathological events leading to severe astrogliosis, glial barrier formation, collagen production, fibrotic scar formation, neuronal death, and motor impairment.

### Gene-silencing of astrocytic *Col1α1* prevents astrogliosis, scar formation, neuronal death, and motor impairment

To examine the role of astrocytic COL1 on ECM alteration and pathology in the PT-induced ischemic stroke model, AAV viruses carrying AAV-PHP.eB-pSico-Col1α1-shRNA-GFP were retro-orbitally injected. GFAP-CreERT2 or Aldh1l1-CreERT2 mice, which were validated as astrocyte-specific promotor mice^38,39^, were used for astrocyte-specific gene-silencing of *Col1α1* (Figures 4A and B). For experimental control for shCol1α1, scrambled shRNA with the same form of virus (AAV-PHP.eB-pSico-Scrambled-shRNA-GFP) was used (Figure 4A). At POD28, *Col1α1*-shRNA infected GFAP-CreERT2 mouse brain showed almost complete disappearance of PT-induced COL1 expression, astrogliosis, fibrotic scar, and neuronal loss compared to control Sc-shRNA brain (Figures 4C and S2D). Sholl analysis and quantitative comparisons of GFAP-positive astrocytes revealed a significant reduction in the number of intersections, volume, length, branch depth, and points in the *Col1α1*-shRNA group compared to the control Sc-shRNA group (Figures 4D-I), indicating a significantly reduced morphological complexity. The scar volume was also reduced considerably in the *Col1α1*-shRNA group (Figures 4J and K). Moreover, the PT-induced motor impairment was recovered significantly in the *Col1α1*-shRNA group compared to the Sc-shRNA group (Figure 4L). Similar results were obtained from another astrocyte-specific promoter mouse, Aldh1l1-CreERT2 (Figure 4L). These results indicate that astrocytic COL1 is necessary for PT-induced glial barrier, fibrotic scar, neuronal death, and motor impairment.

**Figure 4.**
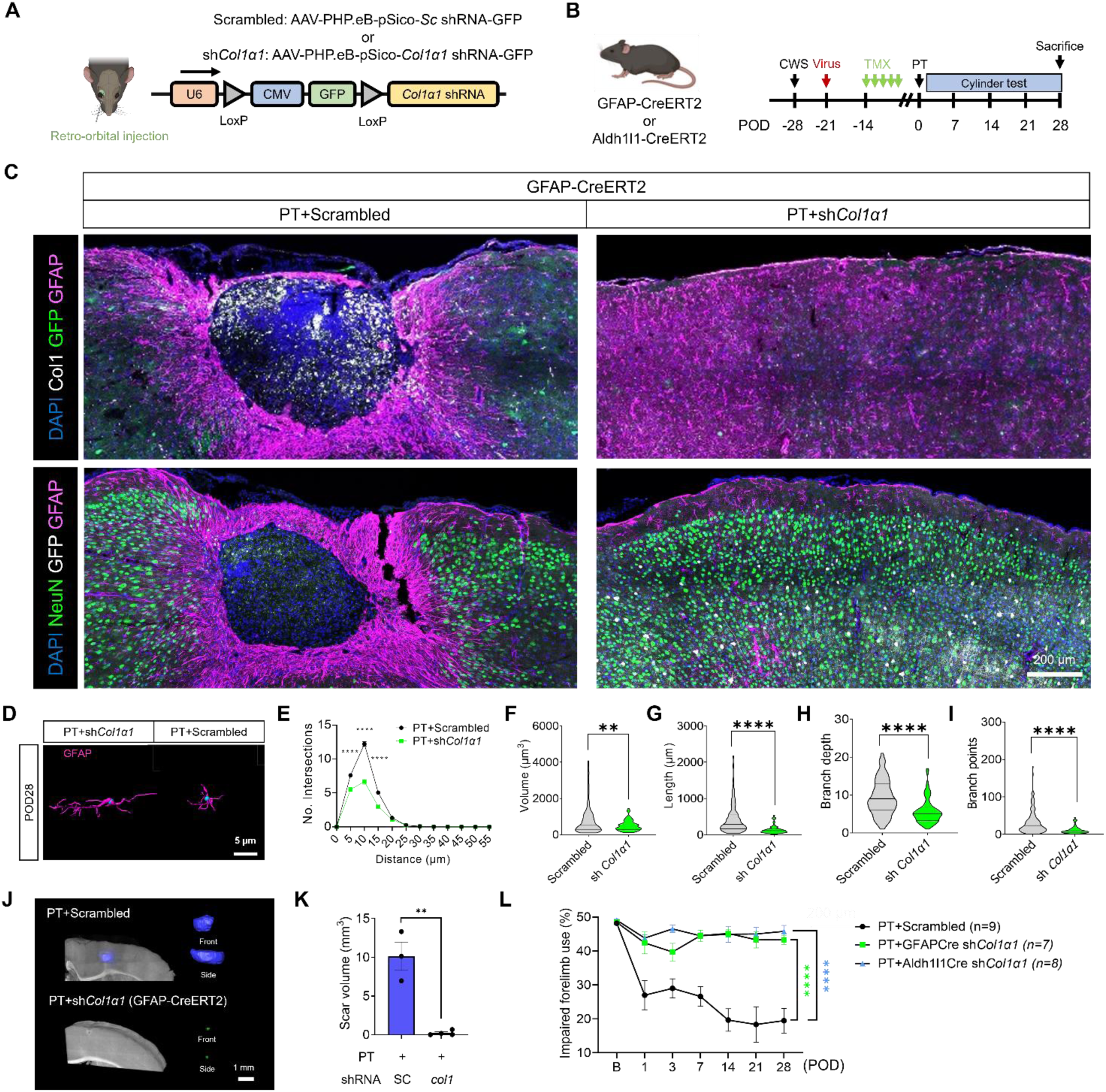
Gene-silencing of astrocytic COL1 prevents astrogliosis, scar formation, neuronal death, and motor impairment. (A) Schematic diagram of *Col1α1* shRNA virus construct with injection route (B) Mouse information with experimental schedule (TMX: tamoxifen injection) (C) Representative images of infarction region in Scrambled and shCol1α1 groups at POD28 (D) Reconstructed image of GFAP signal in PT+Scrambled and PT+sh*Col1α1* groups at POD28 (E) Sholl analysis of astrocytes in PT+Scrambled and PT+shCol1α1 groups (Two-way ANOVA with Bonferroni post hoc, \*\*\*\**p*<0.0001) (F-I) Quantification of Volume (F), Length (G), Branch depth (H), and Branch points (I) in PT+Scrambled and PT+sh*Col1α1* group at POD28 (Unpaired t test, \*\**p*<0.01, \*\*\*\**p*<0.0001) (J) SOCM image and reconstructed scar in PT+Scrambled and PT+shCol1a1 groups (K) Quantification of scar size in PT+Scrambled and PT+sh*Col1α1* groups (error bar: SEM, Mann-Whitney’s U test, \**p*<0.05) (L) Ratio of impaired forelimb used in PT+Scrambled and PT+sh*Col1α1* in GFAP- and Aldh1l1-Cre groups (error bar: SEM, B: baseline, POD: post operation day, Two-way ANOVA with Bonferroni post hoc, \*\*\**p*<0.001, \*\*\*\**p*<0.0001)

### KDS12025, an H_2_O_2_-decomposing peroxidase enhancer, reverses the PT-induced cascade of pathological events

To test whether H_2_O_2_is necessary for astrocytic COL1 production, we treated the PT-induced ischemic stroke mouse model with KDS12025 to reduce aberrant H_2_O_2_levels. KDS12025, a newly developed blood-brain barrier-permeable enhancer of hemoglobin’s H_2_O_2_-decomposing peroxidase activity^40^, was administered via intraperitoneal (IP) injection during the golden time, 5 minutes after PT on the first day, followed by a once-daily dose for two consecutive days (0.1 mg/kg/day for a total of three times; Figure 5A). At POD28, KDS12025 treatment significantly reduced scar volume compared to the PT+vehicle group, as revealed by SOCM (Figures 5B and C). We observed a markedly smaller formation of the glial barrier and profoundly less expression of COL1 in the PT+KDS12025 group compared to the PT+vehicle group (Figures 5D and S8). Detailed morphological analyses showed that astrocytic hypertrophy was significantly reduced in the PT+KDS12025 group (Figures 5E-J). The results from the cylinder test showed significant and dose-dependent recovery from motor impairment in the PT+KDS12025 groups with a highly potent half-maximal inhibitory concentration (IC_50_) of 0.0027 mg/kg/day (Figure 5K).

**Figure 5.**
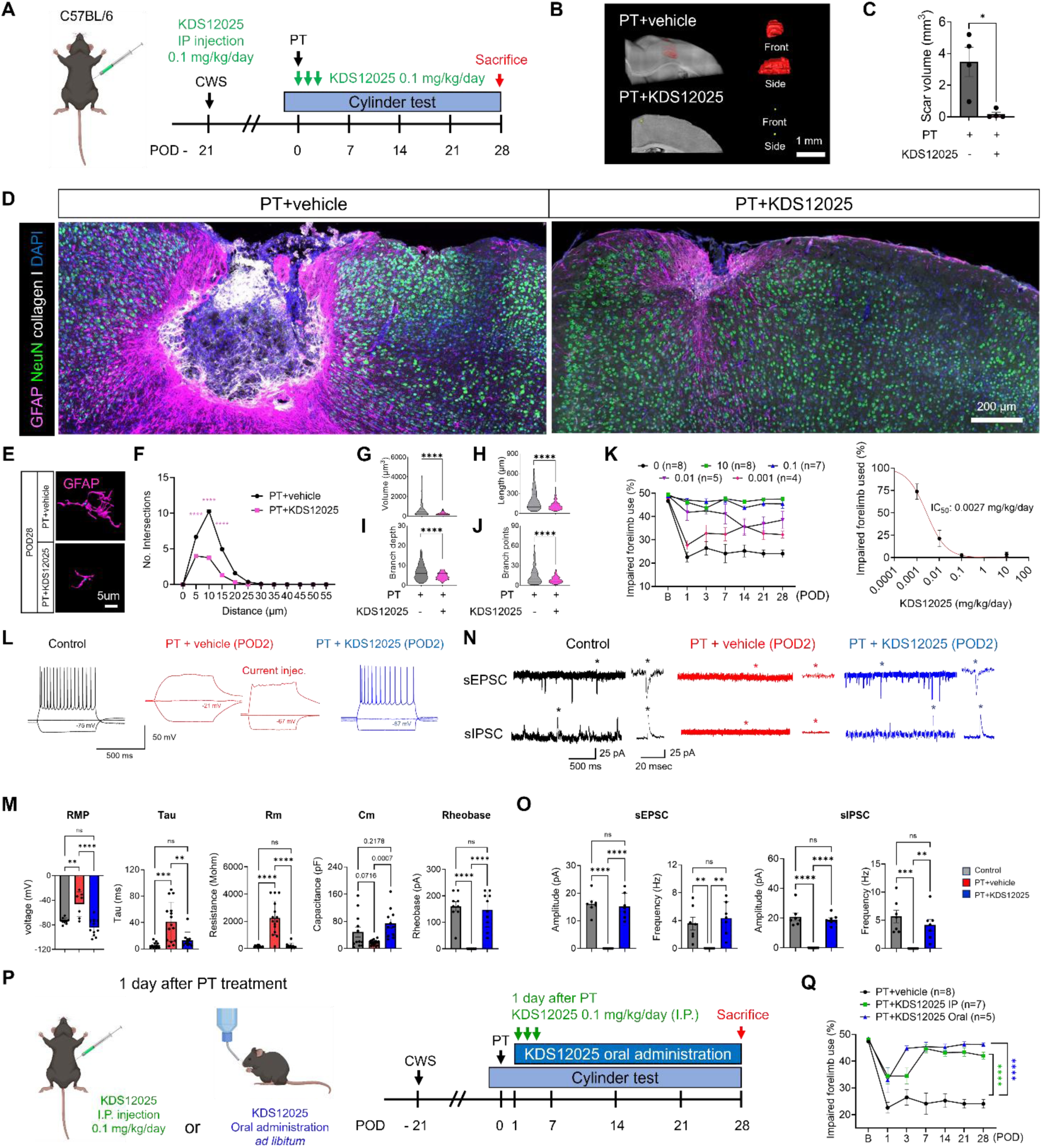
KDS12025, an H_2_O_2_-decomposing peroxidase enhancer, reverses the PT-induced cascade of pathological events. (A) Timeline of the experimental procedure and drug treatment route (CWS: cranial window surgery, PT: Photothrombosis) (B) SOCM image and 3D reconstruction of fibrotic scar in PT+vehicle and PT+KDS12025 groups (C) Quantification of scar size in PT+vehicle and PT+KDS12025 groups (error bar: SEM, Mann-Whitney U test, \**p*<0.05) (D) Representative image of DAPI (blue), collagen I (white), NeuN (green), and GFAP (magenta) expressions at scar regions in PT+vehicle and PT+KDS12025 group at POD28 (E) Representative reconstructed GFAP signal in PT+vehicle and PT+KDS12025 at POD28 (F) Sholl analysis of astrocytes in PT+vehicle and PT+KDS12025 groups (Two-way ANOVA with Bonferroni’s post hoc, \*\*\*\**p*<0.0001) (G-J) Quantitative analysis of astrocyte morphology; dendrite volume (G), dendrite length (H), branch depth (I), and branch point (J) (Unpaired *t* test, \*\*\*\**p*<0.0001) (K) Ratio of impaired forelimb use in various KDS12025 treatment groups (B: baseline, POD: post operation day, error bar: SEM) and dose response curve of KDS12025 at POD28 (error bar: SEM) and dose response inhibition curve of KDS12025 (L) Represent traces after current injection in Control, PT+vehicle, and PT+KDS12025 groups (number within traces indicate membrane potential) (M) Quantification of RMP, Tau, Rm, Cm, and Rheobases in Control, PT+vehicle, and PT+KDS12025 groups (One-way ANOVA with Bonferroni’s post hoc, \*\**p*<0.01, \*\*\**p*<0.001, \*\*\*\**p*<0.0001) (N) Representative EPCSs and IPSCs in Control, PT+vehicle, and PT+KDS12025 groups (magnified traces from asterisks) (O) Quantification of amplitudes and frequencyes of EPSCs and IPSCs in Control, PT+vehicle, and PT+KDS12025 groups (One-way ANOVA with Bonferroni’s post hoc, \*\**p*<0.01, \*\*\**p*<0.001, \*\*\*\**p*<0.0001) (P) Timeline of experimental procedure and drug treatment route (CWS: cranial window surgery, PT: photothrombosis) (Q) Ratio of impaired forelimb use in both KDS12025 treatment groups (B: baseline, POD: post operation day, error bar: SEM, Two-way ANOVA with Dunnett’s post hoc, *p<0.05, ****p<0.0001). Both oral and IP injection of KDS12025 at one day after PT showed significant functional recovery

NeuN positive signals remained at POD3 (Figure 3D and 3G), far beyond the golden time. However, whether these NeuN-positive cells are viable, functional, or recoverable remains unknown. To determine their physiological status and potential for recovery, we performed the whole-cell patch-clamp on neurons from the infarct region of the acutely prepared brain slices at POD2. Our results showed that neurons were still patchable (although much more challenging to patch than the control neurons) (Figure 5L), indicating that the neurons were still alive. However, the neurons of the PT+vehicle group were significantly depolarized with membrane potential (Vm) near -20 mV, significantly impaired membrane properties (time constant Tau, Rm, and Cm), and seriously impaired in their ability to fire action potential upon positive current injections with a complete absence of action potential (Figure 5L and M). Even after adjusting the Vm to around -80 mV by a negative holding current injection, there was no sign of an action potential elicited by a positive current injection (Figure 5L and M), indicating an almost complete absence of the voltage-gated sodium channels. Under voltage-clamp, these neurons displayed completely absent spontaneous excitatory and inhibitory synaptic currents (sEPSC and sIPSC) (Figures 5N and O). In contrast, the patched neurons from the PT+KDS12025 group showed practically complete recovery from the impaired neuronal excitability and synaptic activities induced by PT (Figures 5L-O). We performed immunohistochemistry on the patch-clamped brain slices and observed that the NeuN positive neurons lacked MAP2 signals in the infarct region of the PT+vehicle group, whereas the PT+KDS12025 group exhibited intact MAP2 alongside NeuN positive neurons (Figure S9). These results indicate that neurons at POD2 are still viable after PT, and KDS12025 treatment effectively mitigates neuronal damage induced by infarction, preserving neuronal excitability and synaptic connectivity.

To evaluate whether we could extend the golden time (within 4 hours in mouse^41^), we administered KDS12025 even one day after PT by oral administration *ad libitum* or IP injection (0.1 mg/kg/day three times). As a result, we found a significant functional recovery in both treatment groups (Figures 5P and Q), indicating that KDS12025 exerts protective effects even after one day, when NeuN-positive neurons are still alive and could revive. Therefore, our results raise a possibility of extending the golden time to one day after stroke using KDS12025. Taken together, these results suggest that H_2_O_2_is necessary and acts as an initial trigger for the cascade of pathological events and propose that KDS12025 is an effective and potent therapeutic drug for ischemic stroke.

### PT-induced changes of N-glycan distribution and gene-silencing of astrocytic FUT8 prevents pathological cascade of events

Next, to investigate the potential PTM of COL1-related proteins, possibly via N-linked glycosylation (Figure 1H), we performed matrix-assisted laser desorption/ionization spectrometry imaging (MALDI-MSI) to explore spatial patterns of the N-glycans in mouse PT models of ischemic stroke, including *Col1α1-*shRNA, KDS12025, and the PT-only control group. Three regions of interest, including normal, glial barrier, and fibrotic scar, were depicted on mass spectrometry image based on the adjacent immunohistochemistry image (Figure 6A). Among 64 N-glycans identified based on MALDI-MSI spectral data (Table S1), the top 25 N-glycans were displayed as a hierarchical clustering heatmap (Figure 6A), which clearly distinguished the fibrotic scar areas of the PT group from those in the other two regions. This was further validated through multivariate analysis using partial least squares-discriminant analysis (PLS-DA) (Figure 6B). From the hierarchical clustering heatmap (Figure 6A) and variable importance in projection (VIP) plot (Figure 6C), we identified that N-glycans with increased relative expression in the fibrotic scar region either lacked fucose or had only one fucose, whereas N-glycans with decreased relative expression in the scar area of the PT group typically had two or more fucoses (Figures 6A and C). These differences were also observed when categorizing N-glycans by the number of fucose residues: Complex or hybrid (C/H) type N-glycans with no fucose or one fucose residue showed considerably increased relative abundance in the core region compared to normal, whereas N-glycans with two or more fucose residues significantly decreased (Figures 6D, S10A-P, and Table S1). The representative images and intensity plots demonstrate conspicuous changes in specific N-glycan expression within the fibrotic scar regions (Figures 6E, 6F, and S10A-P). Interestingly, we found that *Col1α1*-shRNA and KDS12025 treatment fully reversed the PT-induced alterations in N-glycan expression levels in both fibrotic scar and glial barrier regions (Figures 6G, 6H, and S11A-R). These findings suggest the potential roles of PTM of ECM-related proteins, particularly those associated with COL1 and H_2_O_2_, with distinctive N-glycosylation patterns in PT-induced glial barrier and fibrotic scar formation.

**Figure 6.**
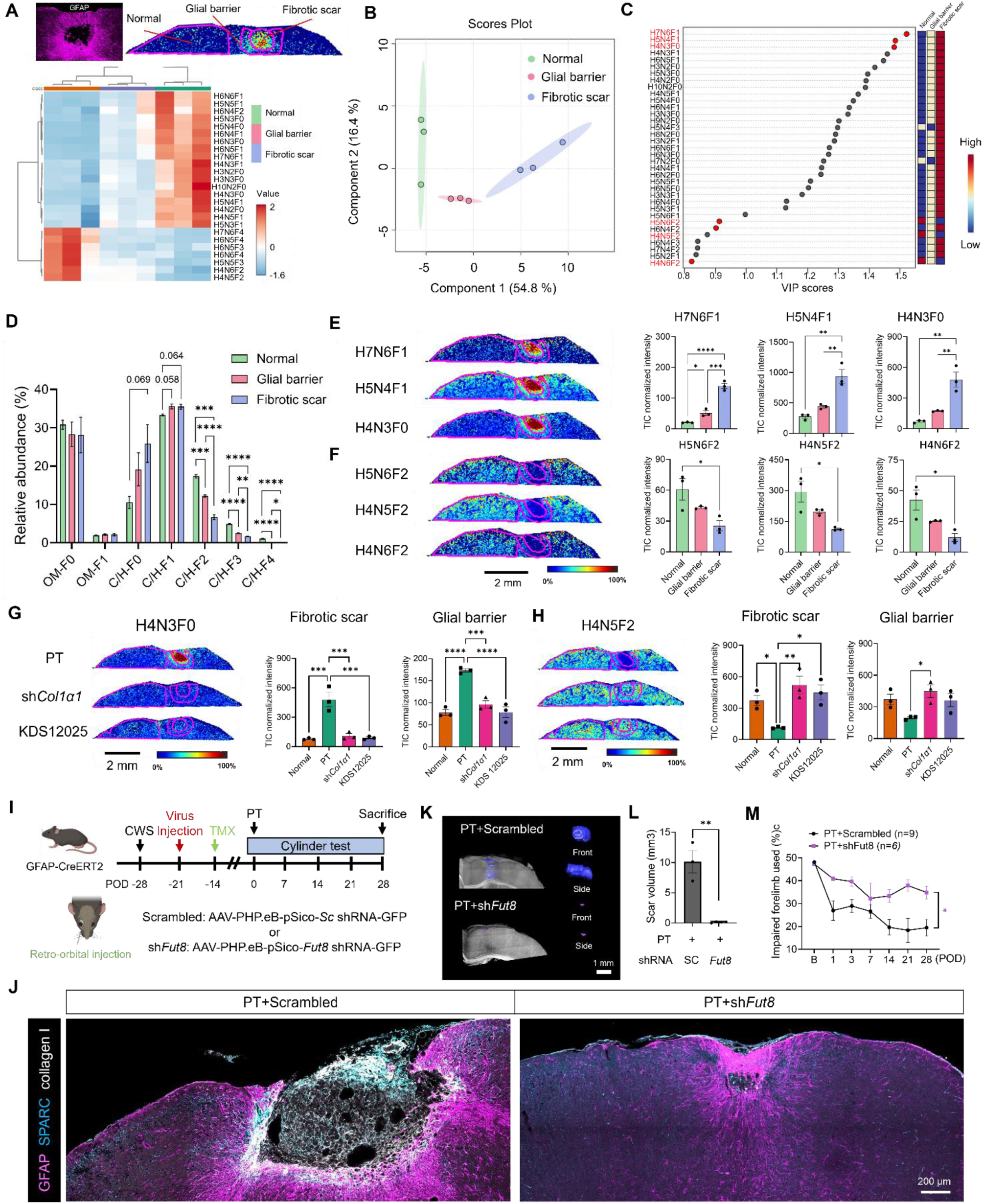
PT-induced changes of N-glycan distribution and gene-silencing of astrocytic FUT8 prevents pathological cascade of events. (A) Representative IHC image for ROI (glial barrier and fibrotic scar) and the hierarchical clustering heatmap of top 25 N-glycans (B) PLS-DA plot of normal, glial barrier, and fibrotic scar (C) VIP score plot of the top 35 N-glycans that contributed to clustering between groups in the PLS-DA plot. The top three N-glycans that increased or decreased in the PT-core region are marked in red (D) Comparison of relative abundance between glycan types in normal, glial barrier, and fibrotic scar (OM: Oligomannose, C/H: complex/hybrid, F0: zero, F1: mono, F2: di, F3: tri, F4: tetra, F5: penta fucosylated glycans, One-way ANOVA with Tukey’s post hoc, \**p*<0.05, \*\**p*<0.01, \*\*\**p*<0.001, \*\*\*\**p*<0.0001), (E and F) The MALDI mass spectrometry images of the six N-glycans (H: Hexose, GlcNAc, N: N-acetylglucosamine, F: fucose, One-way ANOVA with Tukey’s post hoc, \**p*<0.05, \*\**p*<0.01, \*\*\**p*<0.001, \*\*\*\**p*<0.0001) (G and H) Representative MALDI mass spectrometry images and quantification of N-glycan at glial barrier and fibrotic scar regions of PT, sh*Col1α1*, and KDS12025 group (One-way ANOVA with Dunnett’s post hoc, \**p*<0.05, \*\**p*<0.01, \*\*\**p*<0.001, \*\*\*\**p*<0.0001) (I) Timeline of experiment and virus construct information with injection route of *Fut8* shRNA (J) Representative image of infarct region in PT+Scrambled and PT + sh*Fut8* groups (K) SOCM image and 3D reconstructed fibrotic scar (scale bar: 1 mm) (L) Quantification of scar size in PT and PT+sh*Fut8* groups (error bar: SEM, Mann-Whitney’s U test) (M) Ratio of impaired forelimb used in PT+Scrambled and PT+sh*Fut8* (B: baseline, POD: post operation day, error bar: SEM, Two-way ANOVA with Dunnett’s post hoc*p<0.05, **p<0.01)

To investigate further the role of fucosylation in COL1-related proteins, we focused on FUT8 in relation to SPARC. It has been previously reported that silencing of fucosyltransferase 8 (FUT8), the enzyme responsible for core fucosylation, induces conformational changes in SPARC^42^, resulting in reduced COL1 expression and fiber-formation without altering SPARC protein levels in lung stem cells^43^. To provide independent evidence for the role of astrocytic COL1 in the PT-induced pathological cascade, we developed *Fut8*-shRNA and performed astrocyte-specific gene-silencing of FUT8, which could independently modulate astrocytic COL1 production (Figures 6I and S12A-C). At POD 28, we observed an intense expression of SPARC near the inner border of the glial barrier, highly overlapping with COL1 in the Sc-shRNA group, whereas the *Fut8*-shRNA group exhibited almost complete elimination of COL1 and SPARC expression with markedly reduced fibrotic scar and neuronal death (Figures 6J and S12D). Furthermore, SOCM imaging showed almost eliminated scar volume in the *Fut8*-shRNA group (Figures 6K and L). Moreover, the impaired motor function was significantly improved in the *Fut8*-shRNA group (Figure 6M). These results indicate that astrocytic core-fucosylation by FUT8 is also critical for SPARC expression, assembly of collagen fibers, and pathological cascade, proposing astrocytic FUT8 as an effective therapeutic target for ischemic stroke.

### KDS12025 effectively rescues stroke pathology in the PT NHP model of ischemic stroke

As an approved drug for ischemic stroke is currently non-existent, we tested the feasibility of KDS12025 as a small-molecule drug candidate in the NHP model of ischemic stroke. We first employed PT in 3∼4-year-old cynomolgus monkeys (*Macaca fascicularis*) as in the mouse model (Figure 7A). The target infarct area for PT included the dorsal premotor cortex (PMd) and primary motor cortex (M1), which are responsible for hand motor function, especially hand shape and grasp^44,45^ (Figure 7B). KDS12025 was administered orally via esophageal intubation at two different dosage regimens: 1) KDS(0.1); 0.1 mg/kg/day once within 1 hour of golden time and 2) KDS(0.03); 0.03 mg/kg/day within 1 hour after PT on the first day, followed by a once-daily for two consecutive days, total of three times after PT (Figure 7A). A total of three cynomolgus monkeys were used, one for each of three conditions of PT, PT+KDS(0.1), and PT+KDS(0.03). Using magnetic resonance imaging (MRI), we observed that PT-induced brain edema at the infarction site persisted until POD14 in the PT group but was markedly reduced in the PT+KDS(0.1) and PT+KDS(0.03) groups at POD14 (Figures 7C and E). Furthermore, the diffusion tensor imaging (DTI) analysis showed reduced corticospinal tract fibers, which persisted until POD14 in the PT group but markedly recovered in the PT+KDS(0.1) and PT+KDS(0.03) groups at POD14 (Figures 7D and F). To assess motor functions, we performed a single fruit-reaching test before and after PT (Figure 7G). We found that the PT group failed in hand reaching and grasping with a decreased finger grip number (Figures 7G, S14B, and S14C). Strikingly, both PT+KDS(0.1) and PT+KDS(0.03) groups showed remarkable functional recovery in terms of reaching time, success rate, and finger grasp count (Figure 7G and Video S2). These results highlight the high therapeutic potential of KDS12025 as a drug candidate for ischemic stroke.

**Figure 7.**
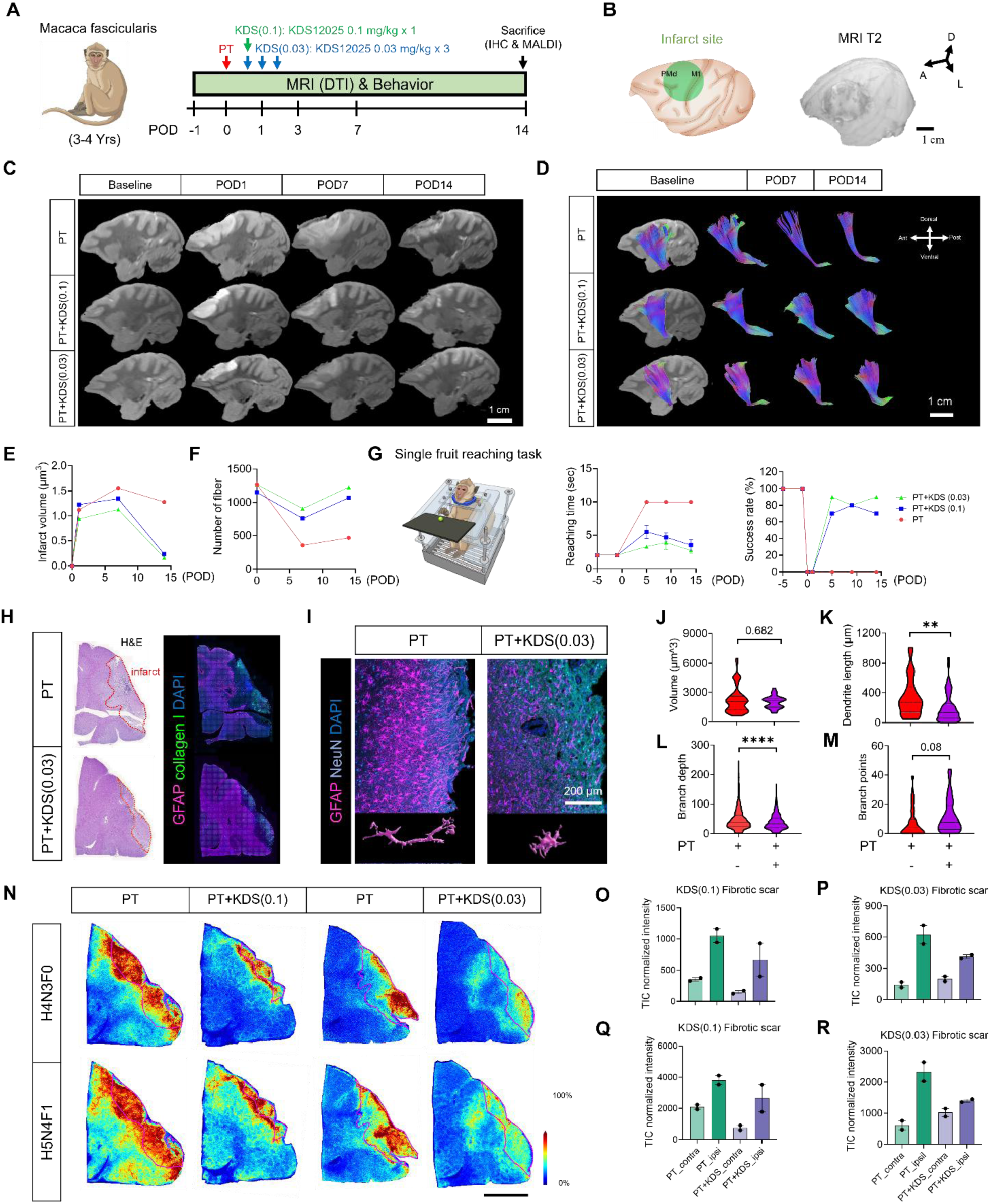
KDS12025 effectively rescues stroke pathology in the PT NHP model of ischemic stroke. (A) Schematic images of experimental schedule (B) Schematic images of target area for PT and MRI T2 image in infarct region at POD1 (PMd: dorsal premotor cortex, M1: primary motor cortex) (C) Serial changes of MRI T2 signals in all experimental groups (D) Diffusion tensor imaging (DTI) at corticospinal pathway in various time points (E) Quantification of infarct volume (F) Quantification of the number of fibers from DTI signal (G) Schematic image of single fruit reaching test and quantification of reaching time (10 seconds limit) and success number (error bar: SEM) (H) Representative H&E and IHC images of scar and glial barrier with GFAP (purple), NeuN (cyan), DAPI (blue) (I) IHC and Reconstructed image of GFAP in glial barrier region (J-M) Quantification of Volume (J), Length (K), Branch depth (L), and Branch points (M) in PT and PT+KDS(0.03) groups (Unpaired *t*-test, \*\**p*<0.01, \*\*\*\**p*<0.0001) (N) Representative MALDI mass spectrometry images of a N-glycan in each group (H: Hexose N: N-acetylglucosamine, and F: fucose) (O-R) N-glycan intensity of H4N3F0 and H5N4F1 in PT+KDS(0.1) and PT+KDS(0.03) groups (error bar: SEM).

To assess the histological features of the infarct, we performed immunohistochemistry and morphological analyses on brain sections of cynomolgus monkeys. We observed COL1-positive signals at the infarct region (Figures 7H, S13, and S14A), as well as GFAP-positive astrocytes at the boundary of the infarct with polarized and hypertrophied morphology in the PT group, which was significantly reduced in the PT+KDS(0.03) group (Figures 7I-M). We also examined the N-glycosylation profile using MALDI-MSI in the scar area of the PT NHP ischemia model. We observed dramatic increases in N-glycan populations, particularly the N-glycans without fucose (e.g., H4N3F0) and with one fucose (e.g., H5N4F1), in the fibrotic scar regions compared to the contralateral regions, which were markedly reduced in PT+KDS(0.1) and PT+KDS(0.03) groups (Figures 7N-R, S15A and S15B). These findings recapitulate the involvement of N-glycosylation in fibrotic scar formation in the NHP ischemia model, just as in the mouse model. Taken together, acute treatment with KDS12025, by preventing excessive H_2_O_2_ accumulation, can effectively reduce astrogliosis, altered N-glycosylation, and infarct volume and preserve motor functions in the NHP ischemic model, highlighting its potential as a therapeutic drug for ischemic stroke in humans.

## Discussion

In this study, we demonstrate for the first time the molecular mechanism for de novo synthesis of astrocytic COL1 and its function in pathological conditions. Transcriptomic analysis with primary culture astrocytes, we found H_2_O_2_ can induce upregulation of genes related to ECM, especially COL1, whose expression is post-transcriptionally disinhibited by miR-29 and post-translationally facilitated by FUT8-mediated fucosylation of ECM proteins such as SPARC. Using the PT-induced ischemic stroke model, the pathological cascade of events starts from an initial surge of H_2_O_2_ near blood vessels, triggering specific astrocytic responses associated with severe reactive astrogliosis marked by an appearance of iNOS and remodeling of the ECM, notably production of COL1. This heightened presence of iNOS and SPARC within astrocytes precedes the secretion and accumulation of COL1. This leads to a glial barrier, ECM remodeling, and fibrotic scar formation, in which the confined neurons are actively killed by toxic COL1 (Figure S16). During these dynamic events, the affected mouse and NHP display severe motor impairment, which persists indefinitely without recovery. In contrast, inhibition of astrocytic COL1 production by either gene-silencing of astrocytic COL1 or FUT8 or pharmacological inhibition of H_2_O_2_ effectively reverses all the symptoms and features of ischemic stroke. Our study unveils unprecedented mechanistic and therapeutic insights that include 1) molecular and cellular mechanism of astrocytic COL1 synthesis, 2) collagen as the killer molecule that slowly exterminates the confined neurons, 3) the identification of H_2_O_2_ as the key triggering molecule and severe reactive astrocytes as the key triggering cell type for the pathological cascade, 4) the existence of a therapeutic window before a glial barrier formation (POD3) during which the NEUN-positive neurons are still present and viable, 5) the actual function of a glial barrier as a confinement for the committed neurons to be killed and a fibrotic scar to replace them, 6) dynamic changes in fucosylation pattern as the critical steps in ECM remodeling, and 7) identification of a series of novel molecular therapeutic targets including astrocytic H_2_O_2_, miR-29, COL1, FUT8, and SPARC to inhibit and almost completely reverse the pathological cascade.

Our study offers the unprecedented role of COL1 as the killer molecule against neurons. COL1 can bind to several forms of integrins^25^, and heterodimers of α1β1 and α2β1 have a high affinity to COL1^46^. Indeed, our findings based on the sandwich culture and cortical injection of H_2_O_2_ experiments corroborate this hypothesis (Figures 3N-P and S8A-H). Using different integrin inhibitors, we found that COL1-induced neuronal deaths were significantly reduced (Figures 2G and H). In support of our data, accumulating lines of evidence suggest an important pathological role of damaged ECM collagens in various neurodegenerative diseases and aging^47^. Based on these findings, we raise a novel concept of “collagenoptosis,” in which COL1 directly and slowly kills neurons. This novel concept of “collagenoptosis” extends the significance of astrocytic COL1 beyond ischemic stroke, providing valuable insights into the pathophysiology of other degenerative diseases, such as amyotrophic lateral sclerosis, retinal degeneration, Parkinson’s disease, and Alzheimer’s disease, which are also associated with excessive ROS production. These exciting possibilities await future investigations.

Different cell types may contribute to COL1 production at both the initial and later stages when scar formation is accompanied by excessive collagen production. Our *in vitro* data show a transient transcriptional upregulation, with increased expression at the initial stage followed by a decrease. Additionally, we found that astrocytic COL1 triggers neuronal death, a phenomenon also observed five days after infarction in an animal PT model. These findings suggest that COL1 produced by severe reactive astrocytes may act as a signaling molecule to initiate the pathological cascade in ischemic stroke associated with excessive H_2_O_2_ production. At later stages, we observe the excessive COL1 expression within the core of the scar region at POD28. However, the source and function of this collagen remain unclear. A recent study suggests that pericyte and perivascular fibroblasts were transcriptionally converged to stromal fibroblast with upregulation of collagen biosynthesis related genes after spinal cord injury^48^. However, the necessity and specific function of this collagen production by stromal fibroblast have not been elucidated. We propose that this excessive COL1 within the core of the scar region may serve as a structural component supporting brain integrity. The reduced compartment caused by neuronal death in the infarct region likely needs to be filled in to stabilize overall brain structure and function. The interplay between the ECM and brain tissue integrity is fundamental for normal function and recovery after injury^49^. The role of ECM components has been suggested in compensating for structural deficits caused by injury, preventing further damage, and facilitating recovery^50^. Future studies investigating the role and functional implications of ECM changes, including type I collagen, are essential to gain a comprehensive understanding of ECM alterations in brain pathology.

Our study provides unprecedented insights into molecular and cellular mechanisms of ischemic stroke. Despite recognizing the critical role of ROS in ischemic stroke, the specific pathway mediating neuronal death has remained enigmatic, often generalized as mitochondrial dysfunction or DNA damage^51,52^. Our study addresses this missing link by elucidating a novel molecular mechanism in which specific levels of H_2_O_2_ induce astrocytic COL1 production, leading to neuronal death. The level of H_2_O_2_ induced by PT is predicted to be more than 200 µM, as COL1 production and neuronal death were not observed at lower concentrations (Figures S3). These results imply that there exists a threshold level of H_2_O_2_ at around 200 µM, which is sufficient to induce the upregulation of various genes related to collagen-containing ECM, including *Col1α1* and *Col1α2*. In other words, if there is a therapeutic approach to keep the H_2_O_2_ level below this threshold, it should prevent the cascade of pathological events leading to neuronal death. Indeed, treatment with KDS12025, a novel H_2_O_2_-decomposing peroxidase enhancer of hemoglobin, effectively lowered the H_2_O_2_ level below the threshold and reversed these pathological symptoms and features in the PT mouse and NHP models even at very low doses. Hemoglobin, predominantly expressed in the blood and endotherial cells, could also be a target of KDS12025 action together with astrocytic hemoglobin^40^. Following a surge of H_2_O_2_ after PT, astrocytes appear to respond immediately as their endfeet oppose the blood vessels. Astrocytes undergo dramatic changes in their cellular properties and morphologies accompanied by molecular switches as evidenced by the initial gradual increase in GFAP, transforming into the barrier-forming severe reactive astrocytes with initial turning-on of iNOS and later turning-on of COL1 and SPARC. Eventually, these GFAP-positive astrocytes transform their morphology into a distinctive radial shape to line up around the fibrotic scar as a building block of the glial barrier. We demonstrate for the first time that this dramatic process requires excessive H_2_O_2_ and COL1 expression. Meanwhile, based on the similar time courses of glial barrier and neuronal death, we hypothesize the actual function of a glial barrier as a confinement for the H_2_O_2_-exposed neurons to be sacrificed and a fibrotic scar to replace them. This idea is strongly supported by the observations that the extent of the glial barrier is highly correlated with the extent of neuronal death and that pharmacological inhibition of H_2_O_2_ and gene-silencing of astrocytic COL1 or FUT8 prevented glial barrier formation and neuronal death. This idea directly contradicts the traditional view that the glial barrier is beneficial and that removing it will exacerbate neuronal death^53^. Future investigations are needed to resolve these two opposing views.

The remarkable protective effect of KDS12025 in both rodent and NHP models offers the potential for treating ischemic stroke. KDS12025 has been recently developed as a highly soluble, blood-brain-barrier-permeable, small molecule, effectively and indirectly reducing H_2_O_2_ by enhancing the pseudoperoxidase activity of hemoglobin^40^. The only clinically available treatment options for ischemic stroke, such as thrombolytic agents and mechanical thrombectomy, often come with significant side effects, including the risk of reperfusion injury. Although targeting the increased ROS levels observed during stroke has emerged as an alternative treatment option for ischemic stroke^54–56^, most direct ROS scavengers have failed to treat ischemic stroke due to multiple confounding factors, such as their high reactivity, impractically high concentration requirements, poor cellular uptake, and low bioavailability^57^. Given that KDS12025 is an indirect H_2_O_2_ scavenger (acting through hemoglobin), the ensured safety, convenient routes of administration (oral or i.p.), and high potency (IC_50_ = 0.0027 mg/kg/day for mouse and 0.03 mg/kg/day for NHP) render KDS12025 as an attractive drug candidate for timely intervention during the golden time (within one hour) of ischemic stroke. Furthermore, by examining the time course of protein expressions, we have identified the existence of an extended therapeutic time window before a glial barrier formation (POD3), during which the NEUN-positive neurons still exist and are viable. Indeed, we found a potent recovery by KDS12025 (0.1 mg/kg/day) even 24 hours after golden time. Future experiments are needed to test if the protective effect of KDS12025 can be extended to POD2 and POD3. Lastly, in addition to the tested three molecular therapeutic targets of H_2_O_2_, astrocytic COL1, and FUT8, we have identified other potential molecular targets for preventing neurodegeneration after ischemic stroke. These include astrocytic iNOS, transiently increased within astrocytes before a glial barrier formation, miR-29, which can degrade *Col1α1* mRNA and switch on COL1 production by reducing its expression, and SPARC, which is essential for COL1 packaging and export into ECM. Future investigations are needed to validate these potential therapeutic targets. Furthermore, because the current study focuses on the acute ischemic stroke model, future explorations of other stroke models, including ischemia/reperfusion and middle cerebral artery occlusion, should follow.

In summary, the newly developed concepts and molecular tools in this study should help further delineate the exact molecular and cellular mechanisms of neuronal death and sculpt effective therapeutic strategies to prevent ischemic stroke-related brain injuries in humans.

## Acknowledgements

We sincerely thank Yejin Cho and Juyeon Chae for their invaluable technical assistance in producing AAV viruses. We also wish to thank the Research Solution Center (RSC) at the Institute for Basic Science (IBS) for managing the animal facilities and providing access to the Imaris software. Special thanks to Dr. Andre Berndt from the University of Washington for generously supplying the oROS-G plasmid, and to Dr. Sohee Yoon from the Korea Research Institute of Standards and Science (KRISS) for sharing the M5 sprayer for MALDI mass spectrometry imaging. Thanks to Hyo-Yeong Oh and Dr. Young-Woong Kim from the Center for Genome Engineering for helping NHP IHC process. Lastly, we are grateful to all members of the C.J.L. laboratory at IBS for their helpful discussions and comments. This study was supported by the Center for Cognition and Sociality (IBS-R001-D2) and through the projects YSF project (IBS-R001-Y2) under the Institute for Basic Science (IBS), Republic of Korea.

## Author contributions

J. L., H. J., I. H., J. L., J. J., M. K., E. L., W. W., E. H. L., W. J., K. P., and S. R. performed the experiments. B. L. and C.J.L. supervised the analysis. J.L. and C.J.L. designed all the experiments and wrote the manuscript with input from the co-authors.

---

**Figure S1.**
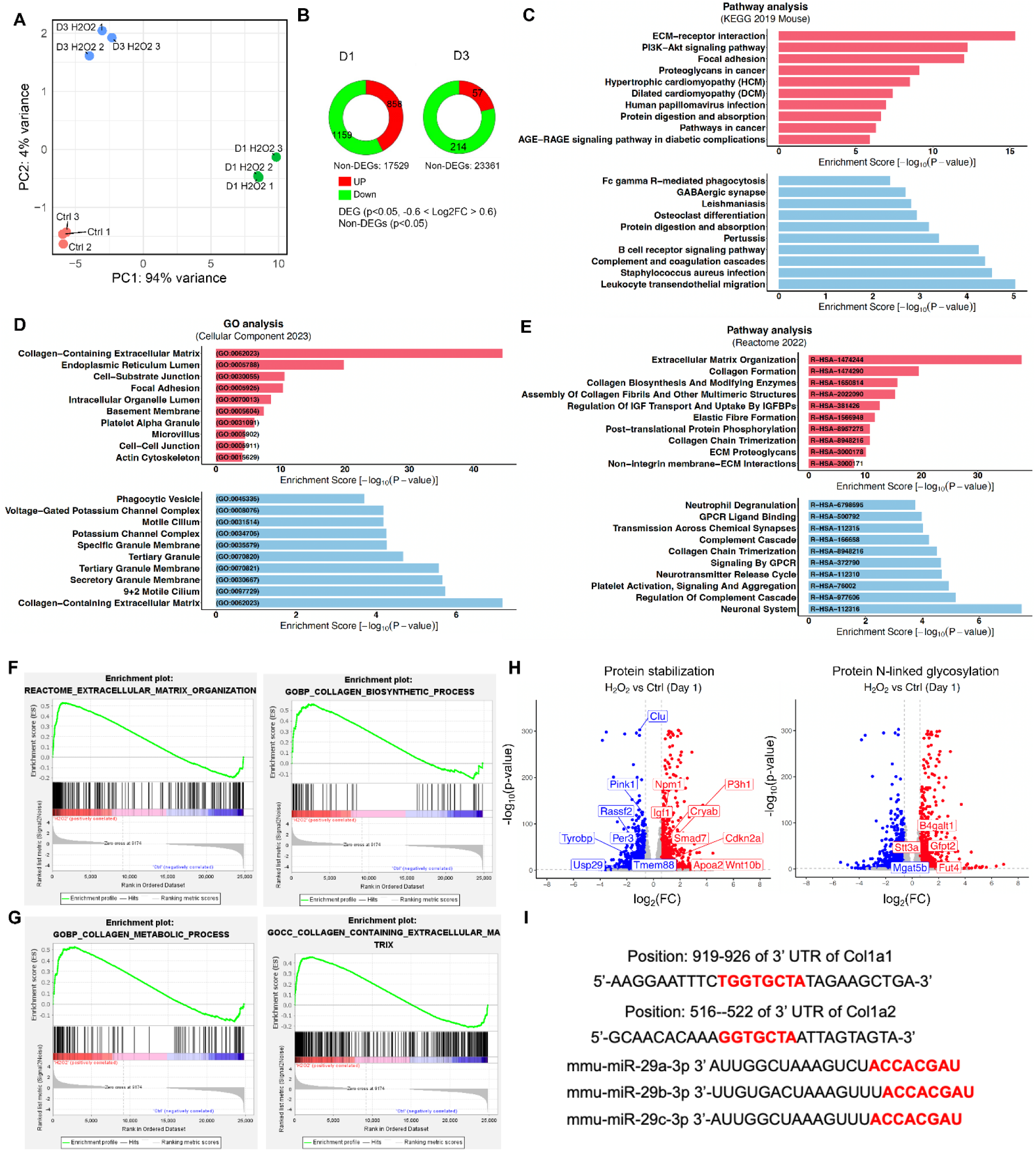
H_2_O_2_ induces collagen-related mRNA changes in astrocyte. (A) PCA plot of RNA-seq samples used. The PCA plot shows the similarities and differences in gene expression patterns among samples. Ctrls (control) and D3s variance (B) Ring plot of total gene numbers with significant *p*-value and DEGs in D1 and 3 (C) Pathway analysis using KEGG database (D) Gene ontology analysis, Cellular Component (E) GSEA analysis of DEGs in D1, Red: Up-regulated genes, Blue: Down regulated genes. The unique GO and Reactome identifiers are shown in the corresponding bar graph (F and G) The enrichment profiles of genes in each terms are shown for Reactome and GO Biological Process (BP) and Cellular Processes (CP) related to collagen. These results indicate H_2_O_2_ induces transcriptional changes, especially related with collagen metabolism and collagen containing ECM (H) Volcano plot of RNA-seq experiment. Gene names of GO:0050821 Protein stabilization and GO:0006487 Protein N-linked glycosylation with *p*<0.05, log2(FC)>|0.6|) in are indicated. Red: Up-Blue: Down-regulation (I) TargetScan prediction results for Col1α1 and Col1α2 in the 3’ UTR and the mouse miR sequences (Red: seed sequence targeted regions)

**Figure S2.**
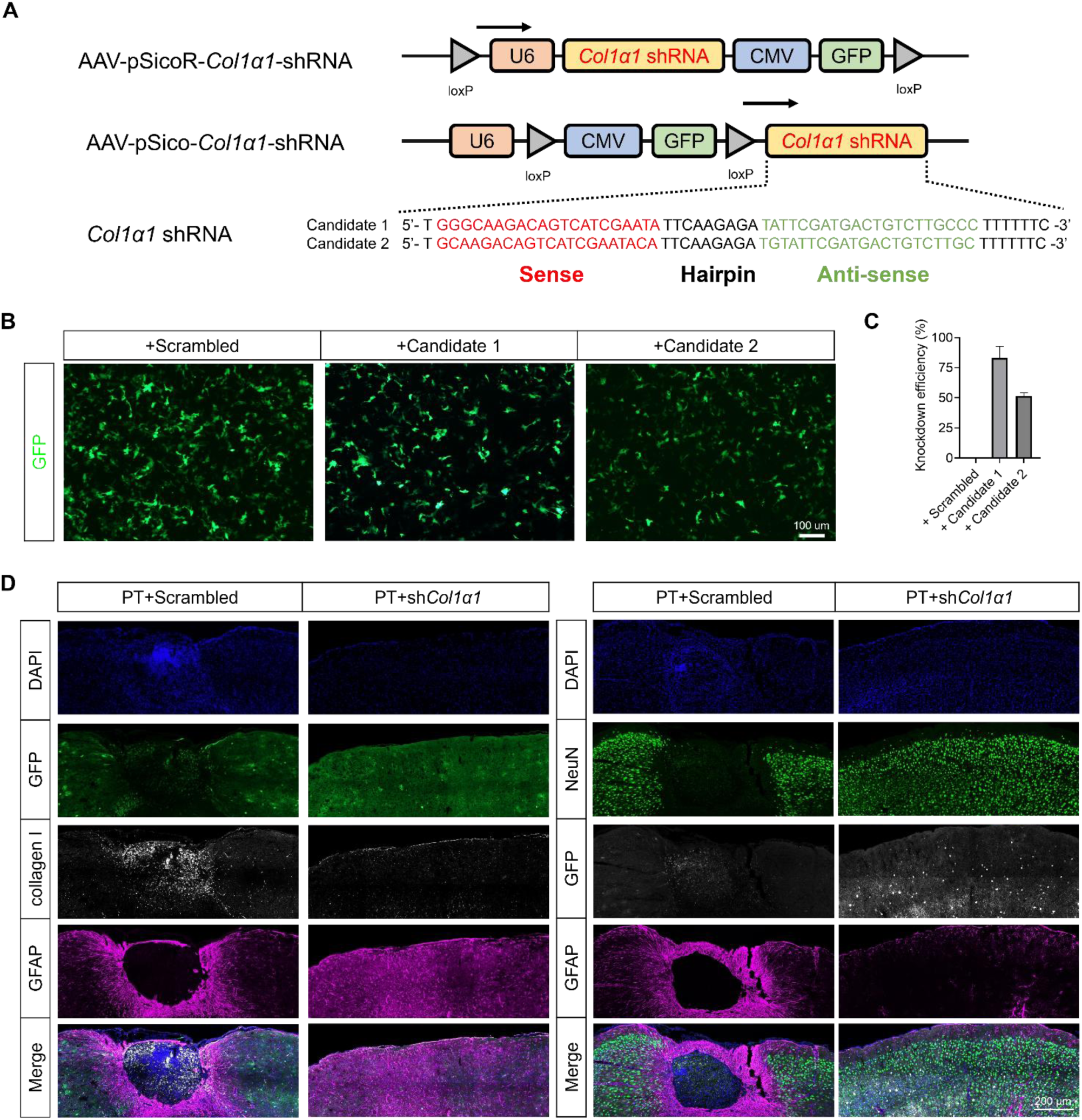
*Col1α1* shRNA effectively reduces *Col1α1* mRNA, and astrocyte-specific gene-silencing of Col1 inhibits collagen production, neuronal death, and glial barrier formation after PT. (A) Schematic virus construct and sequences of *Col1α1* shRNA candidates (B) Representative images of GFP expressions in infected cortical astrocyte (C) Knockdown efficiency of *Col1α1* shRNA candidates, Candidate 1 was selected for both pSico and pSicoR virus package (D) Representative immunostaining images of DAPI (blue), GFP (green), collagen I (white), and GFAP (magenta) in PT+Scrambled and PT+sh*Col1α1* at POD28. Gene-silencing of astrocytic *Col1α1* shows almost no neuronal death and collagen production at the infarct region

**Figure S3.**
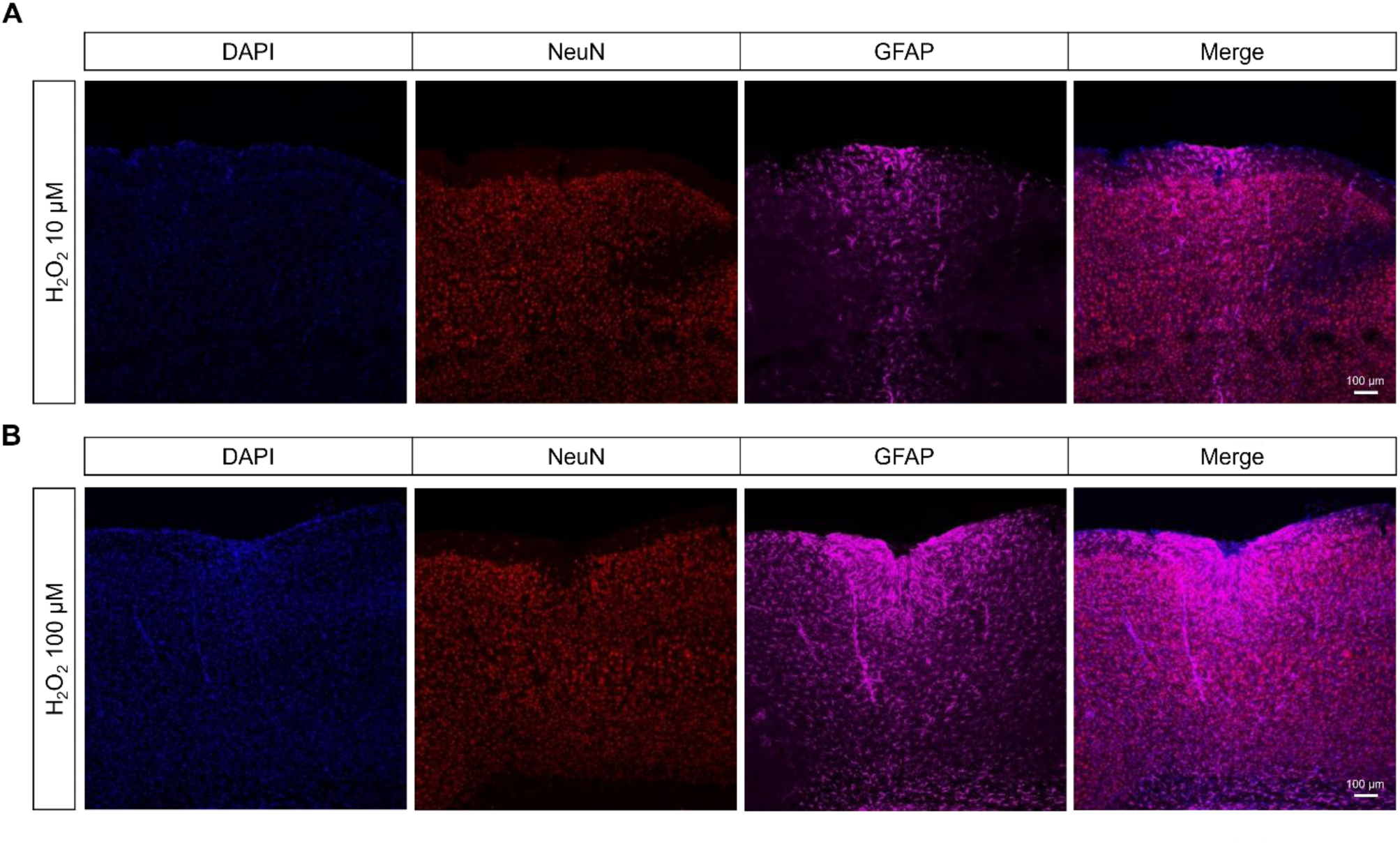
10 and 100 µM H₂O₂ injection induce astrocyte hypertrophy, but not collagen production and neuronal death. (A and B) Representative images of NeuN (red), GFAP (magenta), and DAPI in WT injection with 10 and 100 µM H_2_O_2_ at POD7. H_2_O_2_ injection at 10 and 100 µM did not induce collagen production or neuronal death, indicating that a certain level of H_2_O_2_ is required for astrocytic COL1 production and neuronal death.

**Figure S4.**
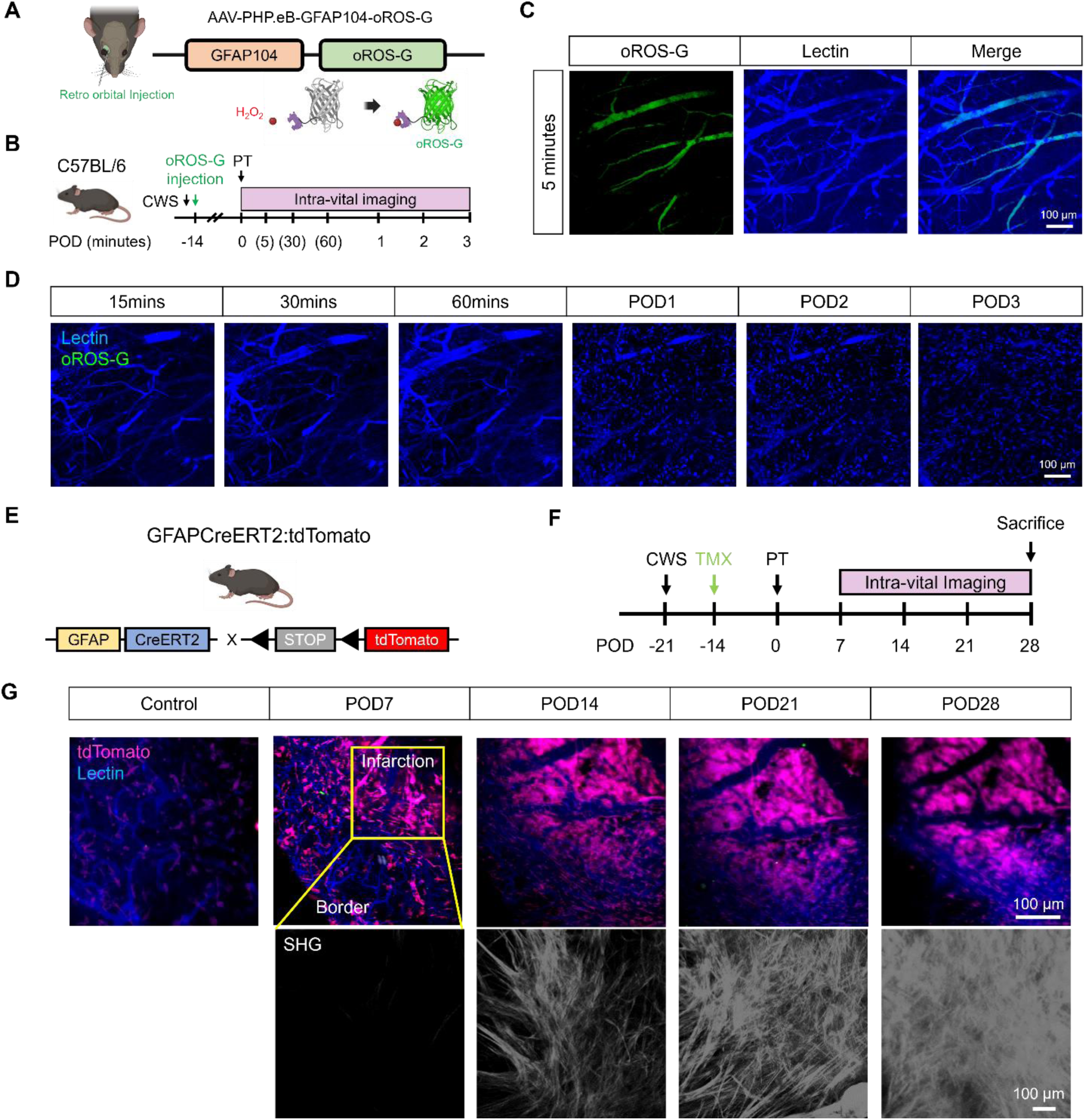
PT causes excessive H_2_O_2_ generation in the initial stage of injury, leading to gradual astrogliosis and collagen production in the infarct region. (A) Virus construct of H_2_O_2_ sensor (oROS-G) with injection route and schematic images of the mode of action (B) Timeline of virus injection and intra-vital imaging (PT: photothrombosis, POD: post operation day) (C) Intra-vital image of Lectin (blue) and oROS-G (green) expressions at 5 minutes after PT (D) Time course images of Lectin (blue) and oROS-G (green) signals, oROS-G signal disappeared from 15 minutes after infarction which implies that disorganization of blood vessel triggers astrocyte end-feet detachment (E) Schematic image of mouse information (F) Timeline of the experiment (CWS: cranial window surgery, TMX: tamoxifen, PT: photothrombosis) (G) Intra-vital images of Lectin (blue), tdTomato (magenta), and SHG (second harmonic generation, white) at infarct region in various time points. Accumulation of astrocyte at the infarct site was observed from POD7 to POD28 while SHC positive signal, indicating collagen fiber was also increased from POD14 to POD28, fully packed at the infarct region

**Figure S5.**
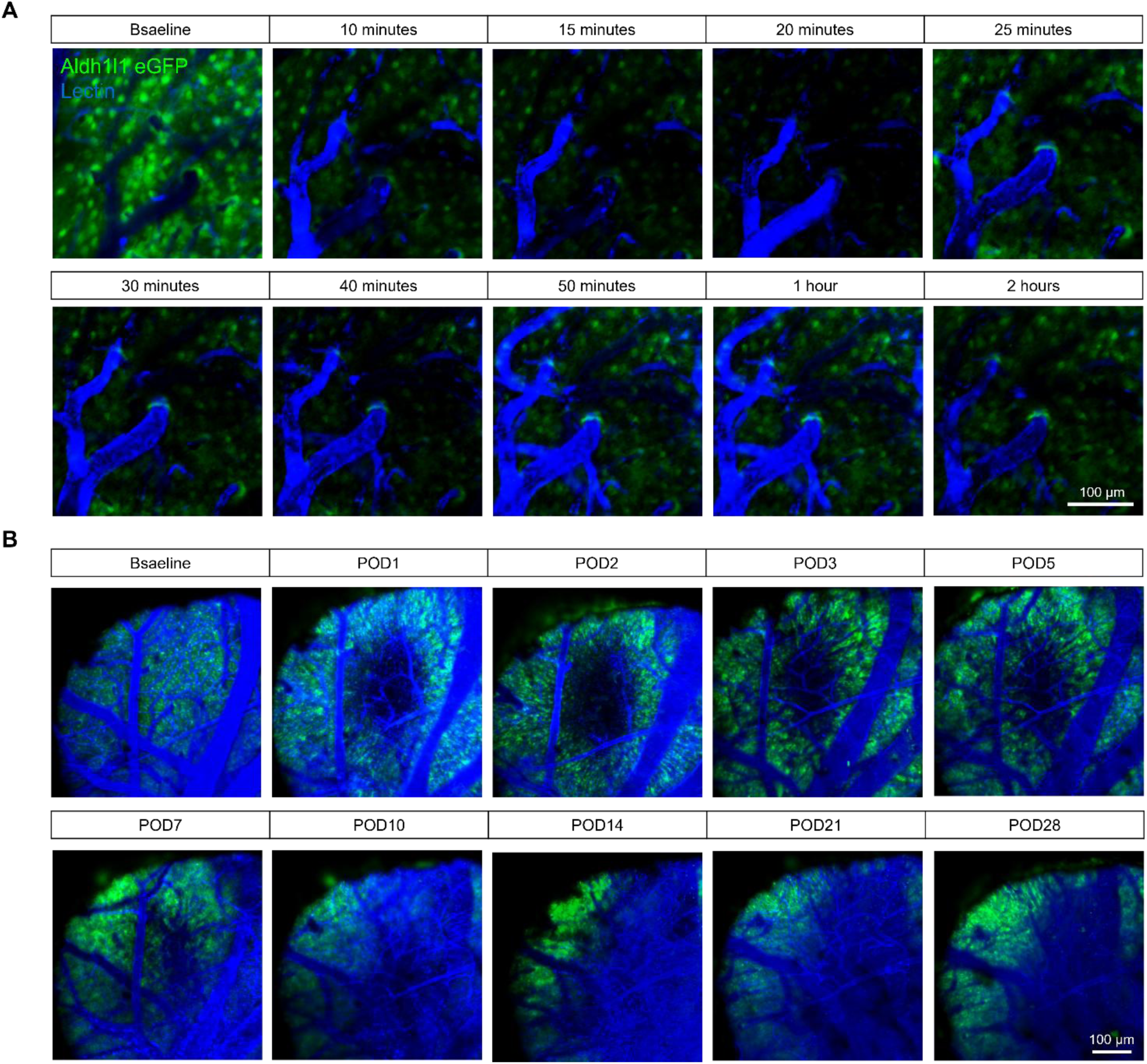
PT causes astrocytic signal changes with dynamic angiogenesis. (A) Changes of astrocyte (eGFP, green) and vascular structure (Lectin, blue) within 2 hours after thrombosis at infarct region (B) Changes of astrocyte and vascular structure until four weeks after thrombosis at the infarct region

**Figure S6.**
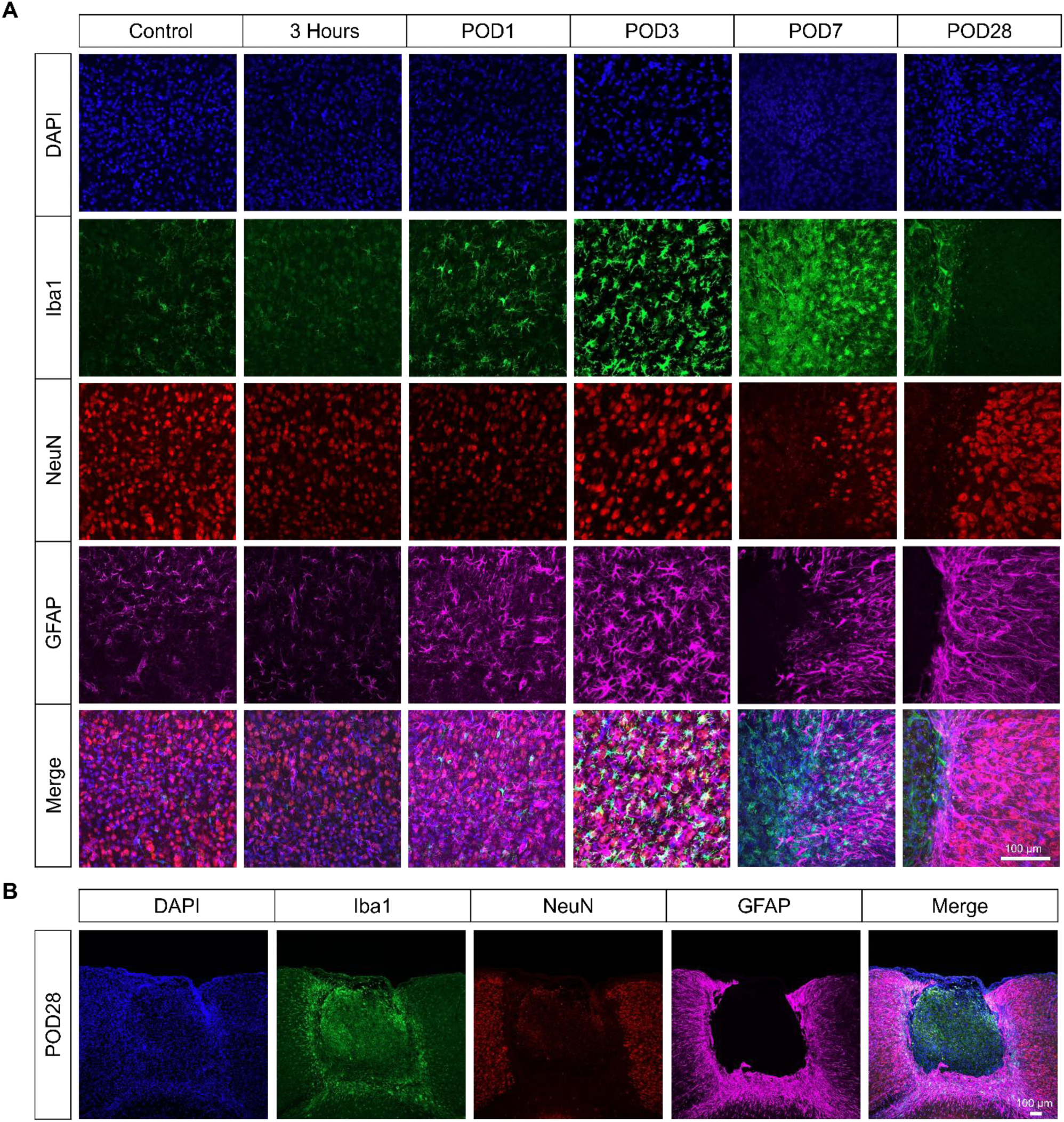
PT causes dynamic expression changes in glial cells and neuron. (A) Time course image of glial cell expressions including microglia (Iba1, green) and astrocyte (GFAP, magenta) and neuron (NeuN, red) in POD3, POD7, and POD28 at infarct region (B) Represent image of microglia (Iba, green), astrocyte (GFAP, magenta), and neuron (NeuN, red) in POD28 at infarct region

**Figure S7.**
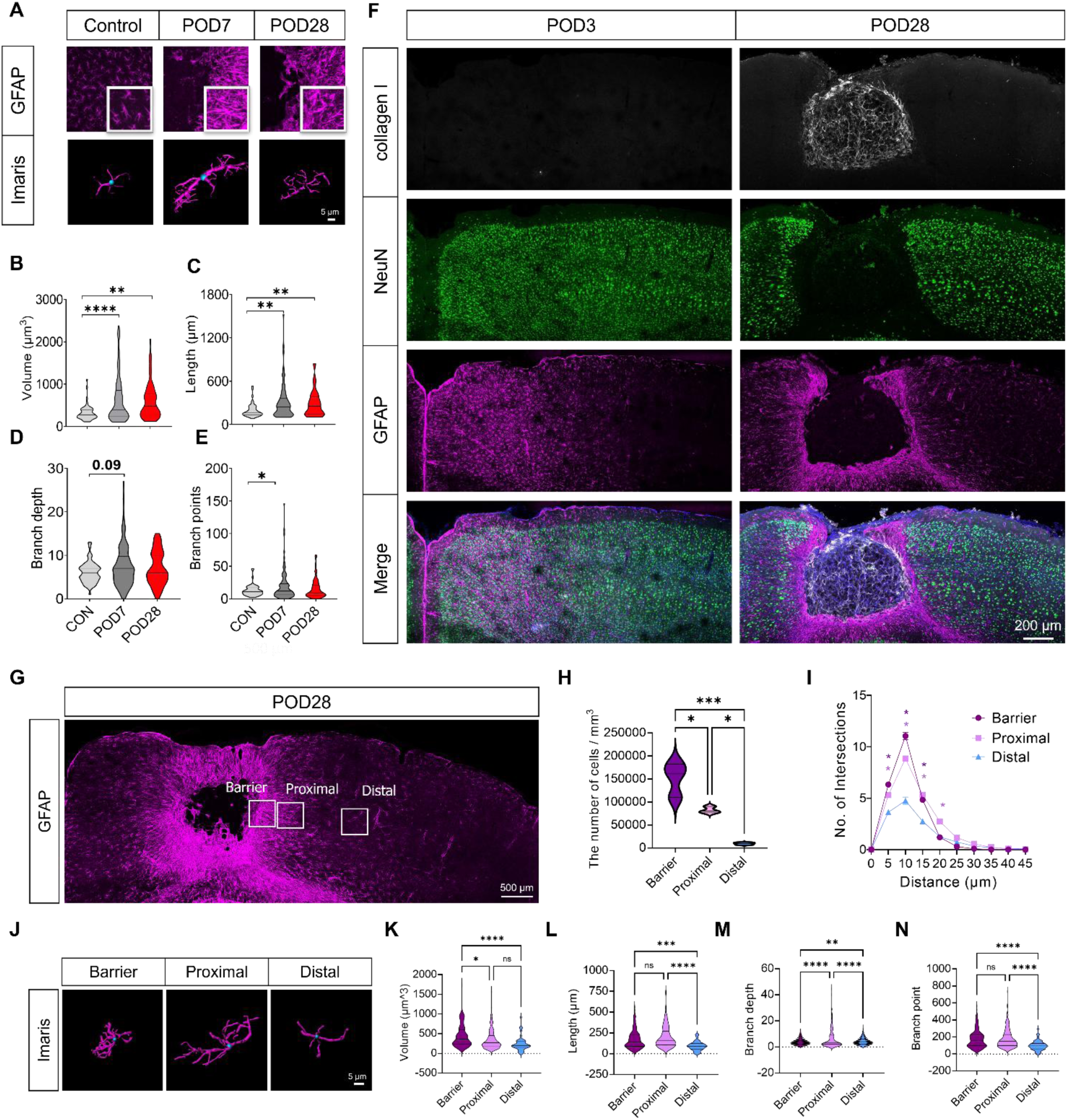
PT induces neuronal death, collagen production, region-specific astrogliosis, and glial barrier formation. (A) Representative GFAP and Imaris reconstructed images in control, POD7, and 28 (B-E) Quantificational analysis of volume (B), length (C), branch depth (D), and branch points (E) at CON (control), POD7, and 28 (One-way ANOVA with Tukey’s post hoc, \**p*<0.05, \*\**p*<0.01, ****p<0.0001) (F) Representative images of collagen I (white), NeuN (green), and GFAP (magenta) at infarct region in POD3 and 28 (G) Representative image of GFAP in barrier, penumbra, and outside regions (H) Number of astrocyte in different regions (One-way ANOVA with Tukey’s post hoc, \*\*\**p*<0.001, \**p*<0.05) (I) Sholl analysis of astrocyte intersections at different regions (Two-way ANOVA with Dunnett’s post hoc, \**p*<0.05) (J) Reconstructed astrocyte morphology at barrier, proximal, and distal regions (K-N) Quantification of volume (K), length (L), branch depth (M), and branch point (N) in astrocytes at different regions (One-way ANOVA with Dunnett’s post hoc, **p<0.01, ***p<0.001, \*\*\*\**p*<0.0001). At POD28, type I collagen packed at fibrotic scar and NeuN signals disappeared at the fibrotic scar and glial barrier region. Astrocytes at the barrier and penumbra regions show significant differences in the number and morphology.

**Figure S8.**
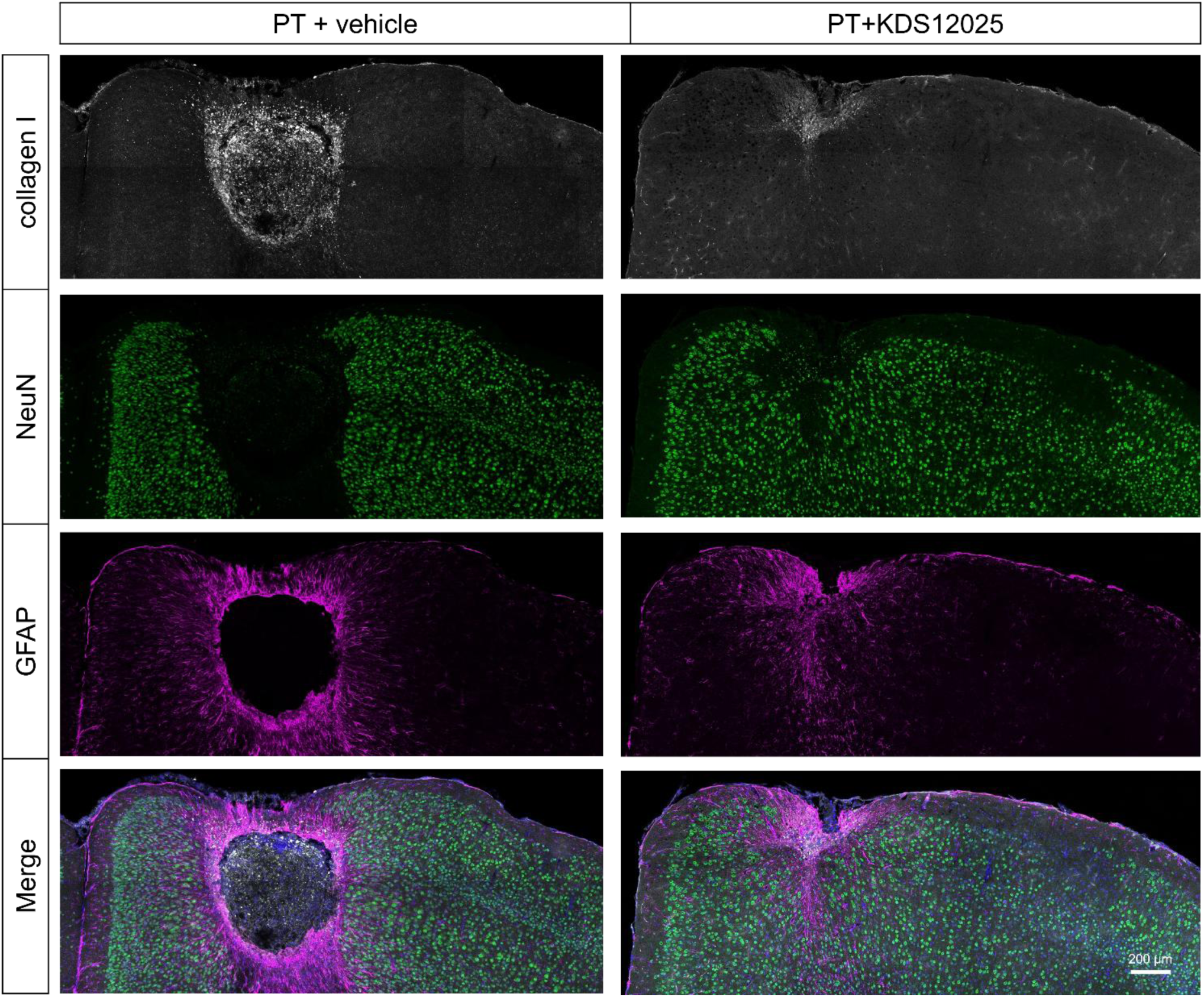
KDS12025 reduces collagen I production and neuronal death at infarct region. Representative image of collagen I (white), NeuN (green), and GFAP (magenta) in infarct region of PT and PT+KDS12025 group at POD28

**Figure S9.**
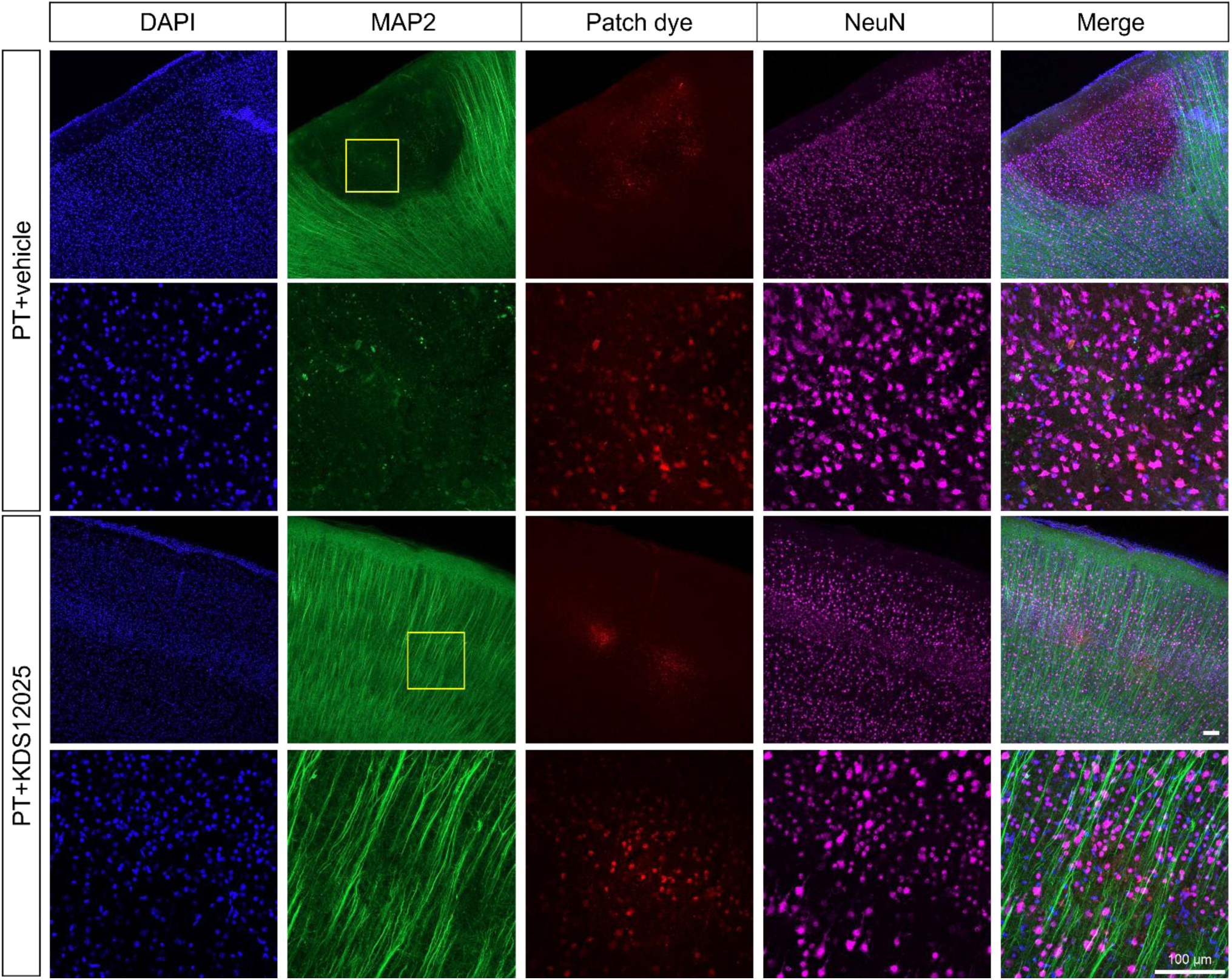
Dendrite degeneration occurred in the PT group at the infarct region while preserved in PT+KDS12025 group. Represent images with a higher magnification of the infarct region of a brain slice after patch-clamp. DAPI (blue), MAP2 (green), Patch dye (alexa 594), and NeuN (magenta)

**Figure S10.**
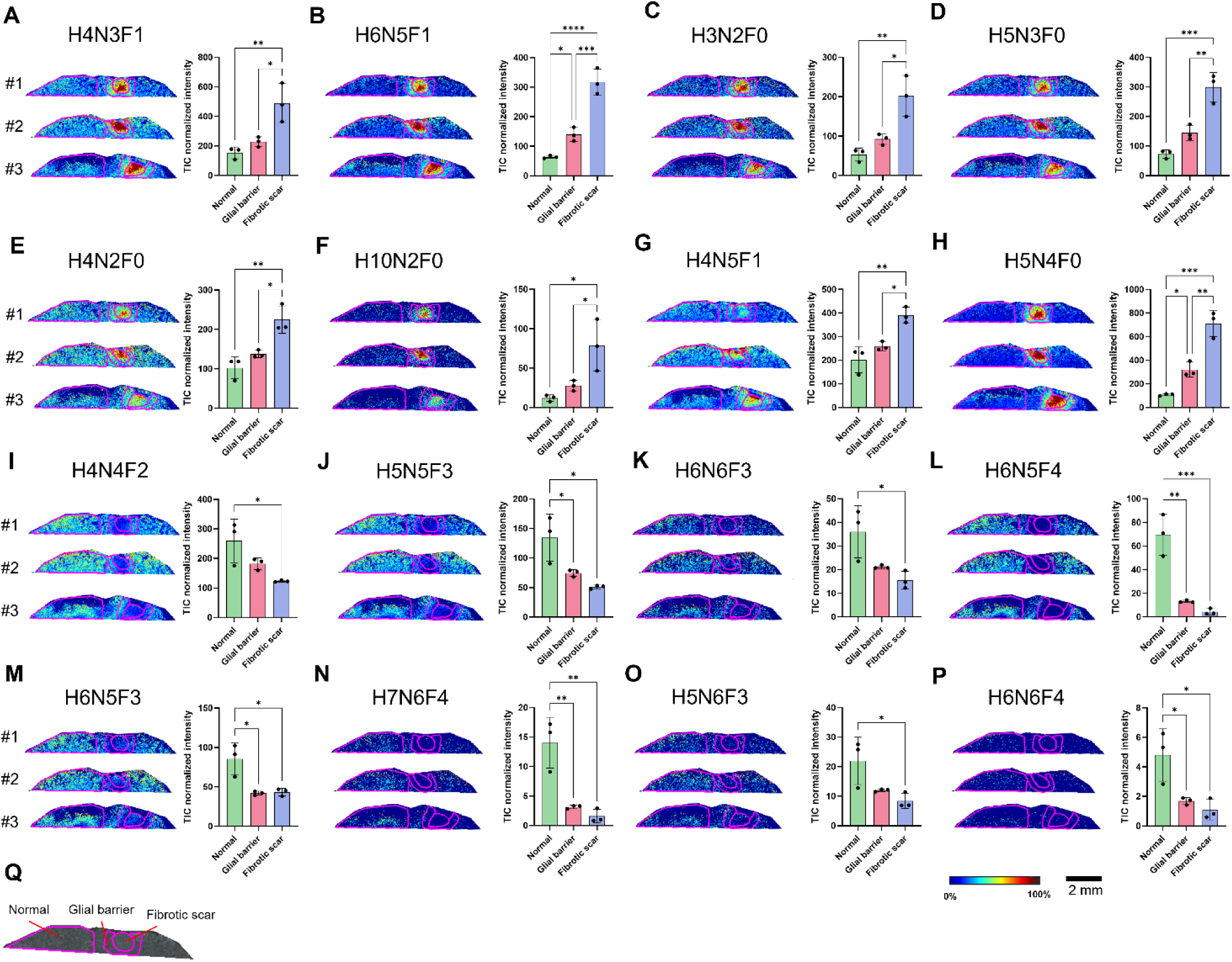
PT induces dramatic changes in fucose-containing N-glycans in a glial barrier and fibrotic scar of the mouse brain. (A-P) MALDI mass spectrometry images and intensity plot of 16 N-glycans (H: Hexose, GlcNAc, N: N-acetylglucosamine, F: fucose) showed significant increases (A-H) or decreases (I-P) in the glial barrier and fibrotic scar (n=3, One-way ANOVA with Tukey’s post hoc analysis. \**p*<0.05, \*\**p*<0.01, \*\*\**p*<0.001, \*\*\*\**p*<0.0001) (Q) Regions of interest (ROIs) include the glial barrier, fibrotic scar, and normal tissue. N-glycans with no fucose or one fucose showed significant increases, whereas those with two or more fucoses showed significant decreases suggesting a possible modulation of the fucosylation pathway by PT.

**Figure S11.**
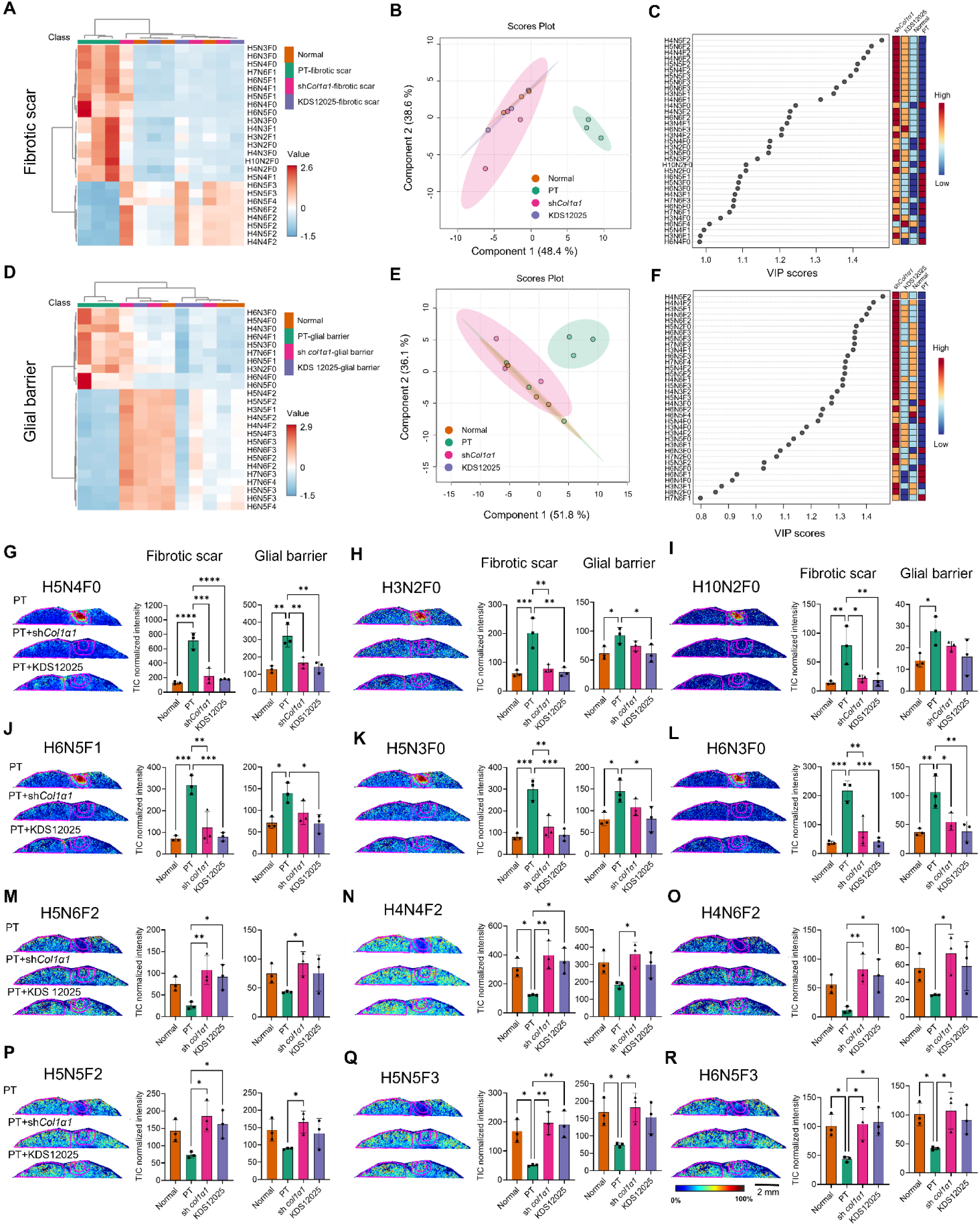
*Col1α1* shRNA and KDS12025 treatment completely reverse the PT-induced changes in fucose containing N-glycans in glial barrier and fibrotic scar regions. (A) Hierarchical clustering heatmap at fibrotic scar region (H: Hexose, GlcNAc, N: N-acetylglucosamine, F: fucose) (B and C) PLS-DA analysis and VIP score at fibrotic scar region in PT, sh*Col1α1*, and KDS12025 group (D) Hierarchical clustering heatmap at glial barrier region (E and F) PLS-DA analysis and VIP score at glial barrier region in PT, sh*Col1α1*, and KDS12025 group (G-R) MALDI mass spectrometry images and the intensity plot of twelve N-glycans showed significant increase or decrease by PT in the fibrotic scar, which was fully restored to normal levels by both sh*Col1α1* and KDS12025 (One-way ANOVA with Dunnett’s post hoc, \**p*<0.05, \*\**p*<0.01, \*\*\**p*<0.001, \*\*\*\**p*<0.0001)

**Figure S12.**
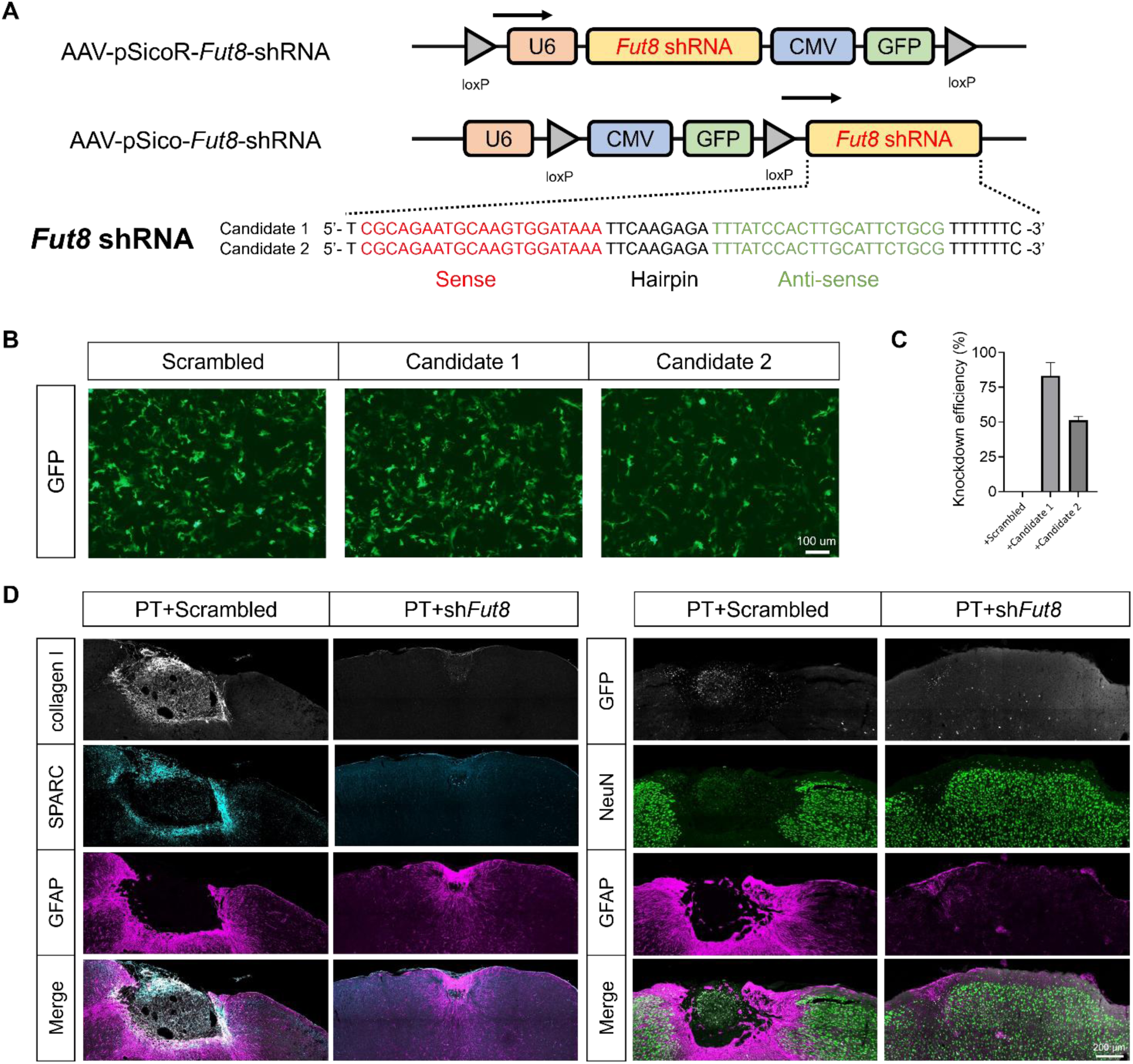
*Fut8* shRNA effectively induces *Fut8* mRNA reduction, and gene-silencing of astrocytic FUT8 reduces fibrotic scar and neuronal death after PT. (A) Schematic virus construct and sequences of *Fut8* shRNA candidates (B) Representative images of GFP expressions in infected cortical astrocyte (C) Knockdown efficiency of *Fut8* shRNA candidates, candidate 1 was chosen for both pSico and pSicoR virus package (D) Representative immunostaining of collagen I (white), SPARC (skyblue), NeuN (green), and GFAP (magenta) in PT+Scrambled and PT+sh*Fut8* at POD28, PT+Scrambled show fibrotic scar and SPARC overlapping collagen I positive signal at the glial barrier region, while PT+sh*Fut8* show reduced fibrotic scar with almost absence of SPARC and collagen I expression at the glial barrier.

**Figure S13.**
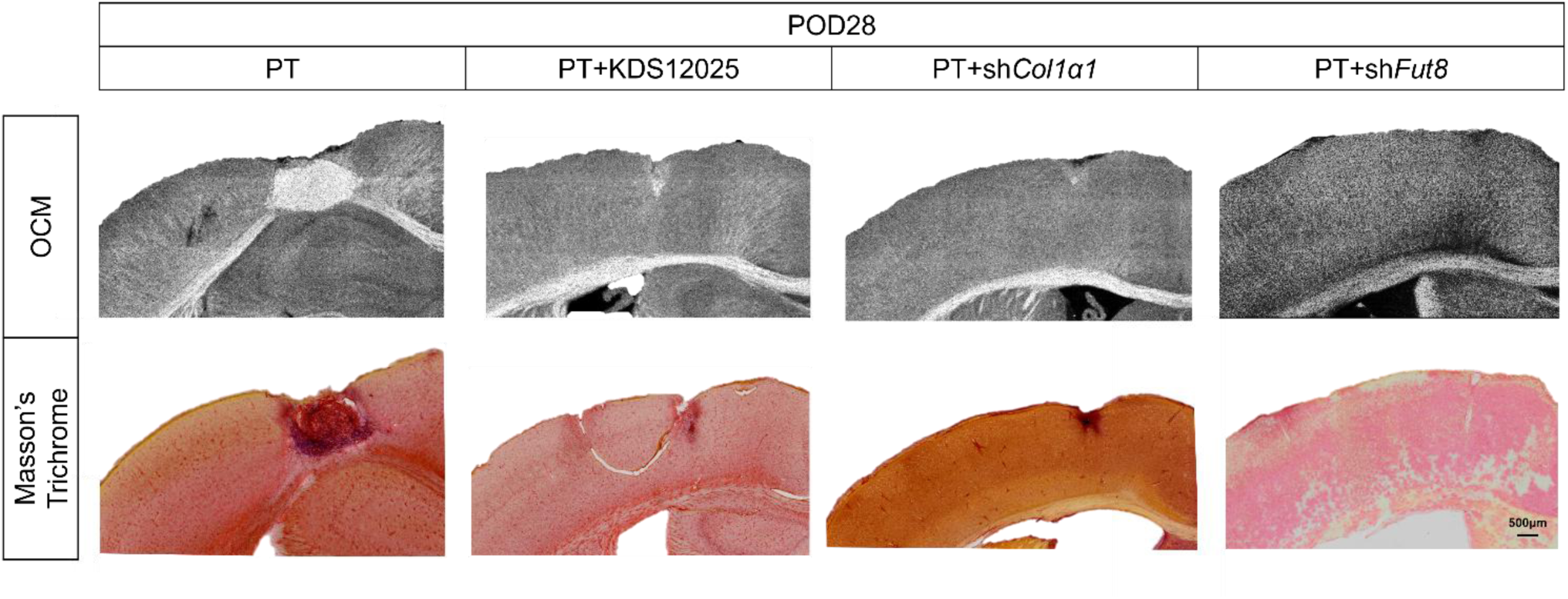
KDS12025, astrocytic COL1, and FUT8 gene-silencing reduce fibrotic scar and collagen production. Representative images of SOCM and Masson’s trichrome staining in PT, PT+KDS12025, PT+sh*Col1α1*, and PT+sh*Fut8*. Red signals in Masson’s Trichrome image represent collagen fiber. Astrocytic gene-silencing of both COL1 and FUT8 resulted in an almost complete absence of fibrotic scar and collagen fiber production.

**Figure S14.**
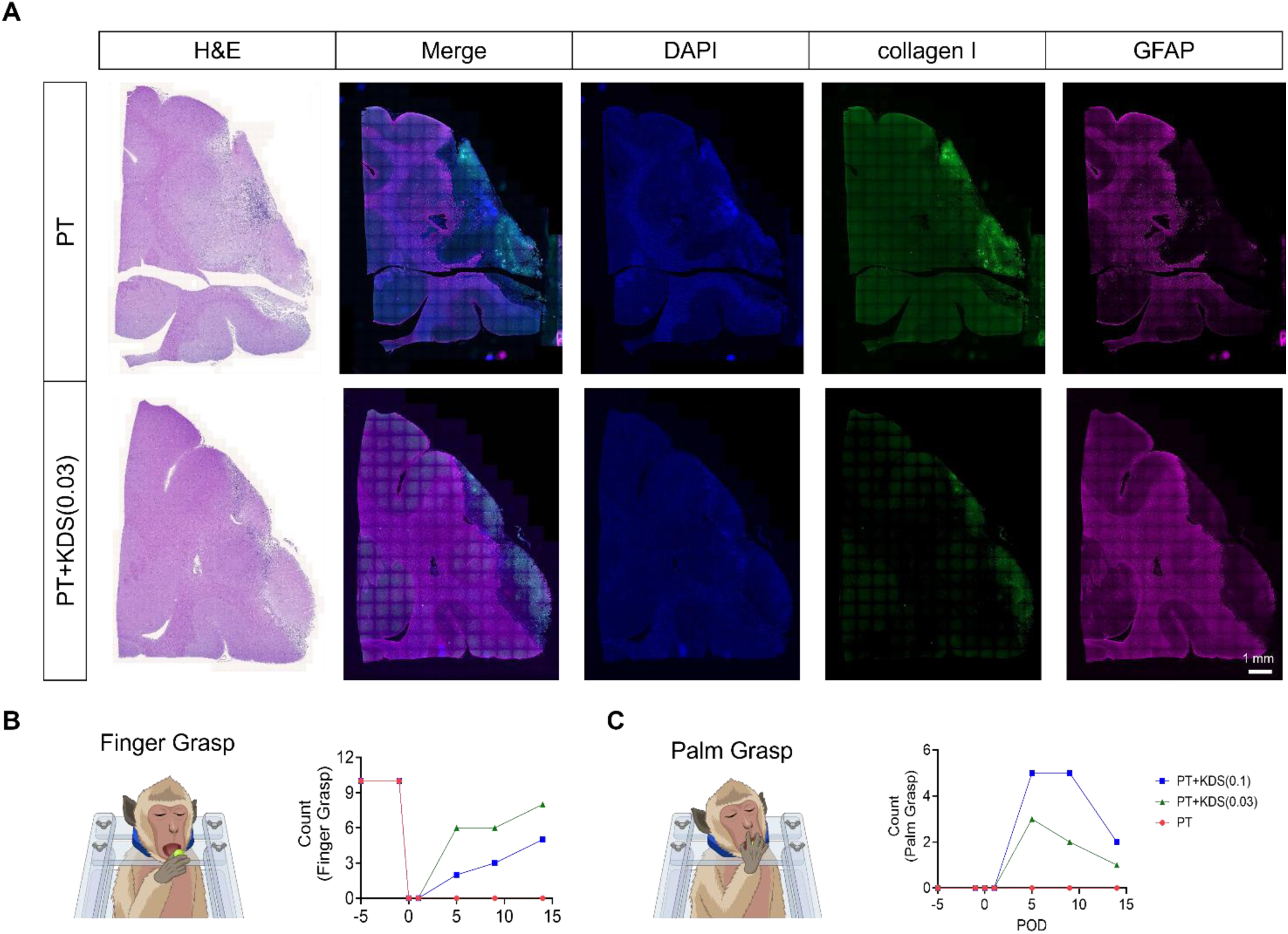
KDS12025 reduces PT-induced collagen production and motor dysfunction in the NHP model of ischemic stroke. (A) Representative images of H&E and immunostaining of collagen I (green) and GFAP (magenta) with DAPI at infarct region in PT and PT+KDS(0.03). This result shows prevention of H_2_O_2_ by KDS12025 reduces COL1 production and fibrotic scar formation (B) Schematic image of finger grasp (using the thumb and fingers to pick up and eat fruit) and quantification of result (C) Schematic image of palm grasp (using the whole hand and palm to push the fruit into the mouth) and quantification of the result. This grasp is typically observed when there is a loss of fine motor control in the fingers. These results indicate that KDS12025 induces fine motor function recovery after ischemic stroke

**Figure S15.**
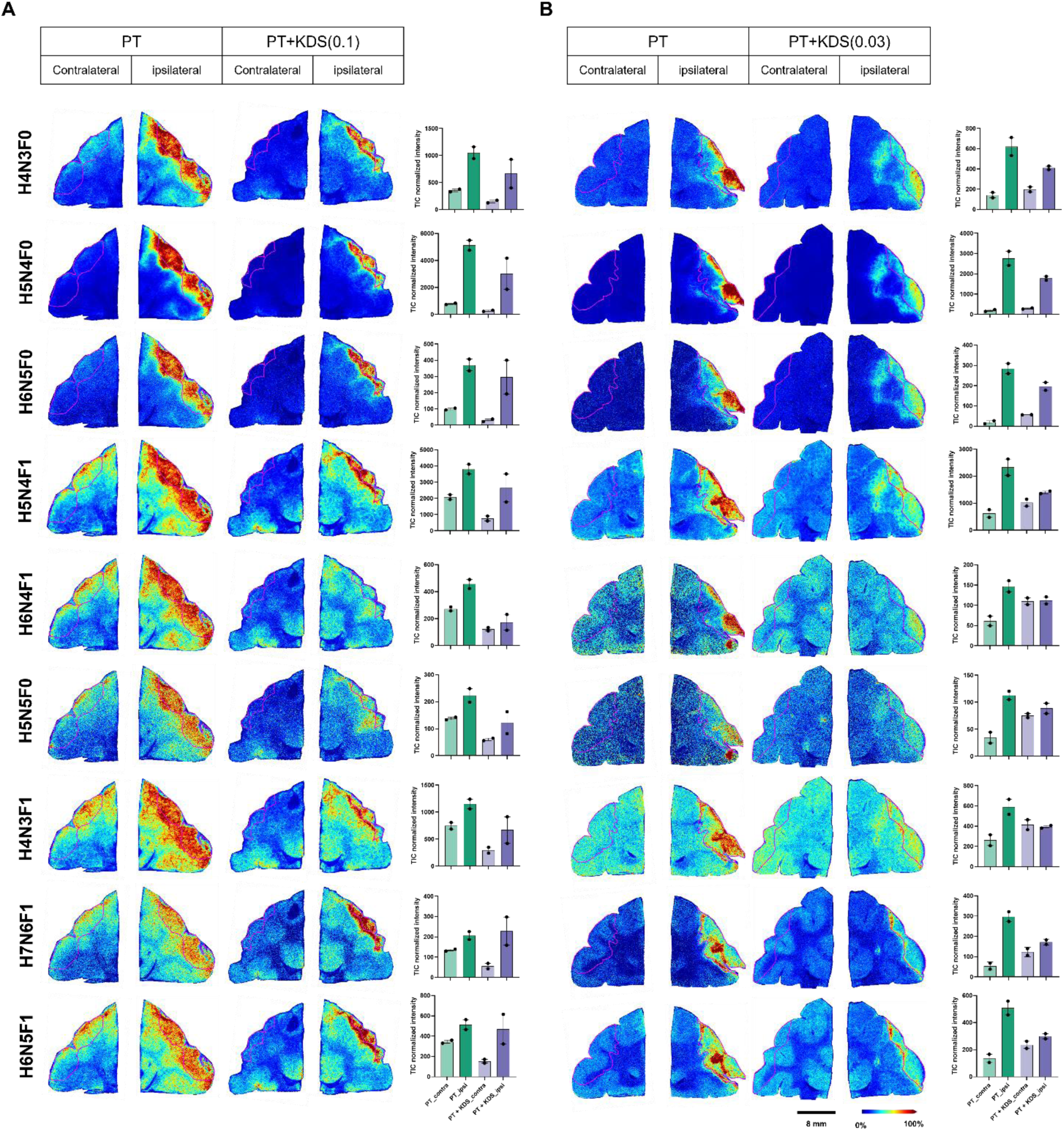
PT induces dramatic changes in fucose containing N-glycans in the infarct regions of PT and PT+KDS12025 treatment groups. (A and B) MALDI mass spectrometry images and intensity plot of nine N-glycans (H: Hexose, GlcNAc, N: N-acetylglucosamine, F: fucose) in the NHP model brains treated with (A) 0.1mg/kg KDS12025 (KDS 0.1) and (B) 0.03mg/kg KDS12025 (KDS 0.03). The bar graph represents the average intensities extracted from two serial sections from one NHP model for each group. These data show that treatment with KDS12025 reduces the increased intensity of N-glycans after PT in an ischemic stroke NHP model. (Error bar: SEM).

**Figure S16.**
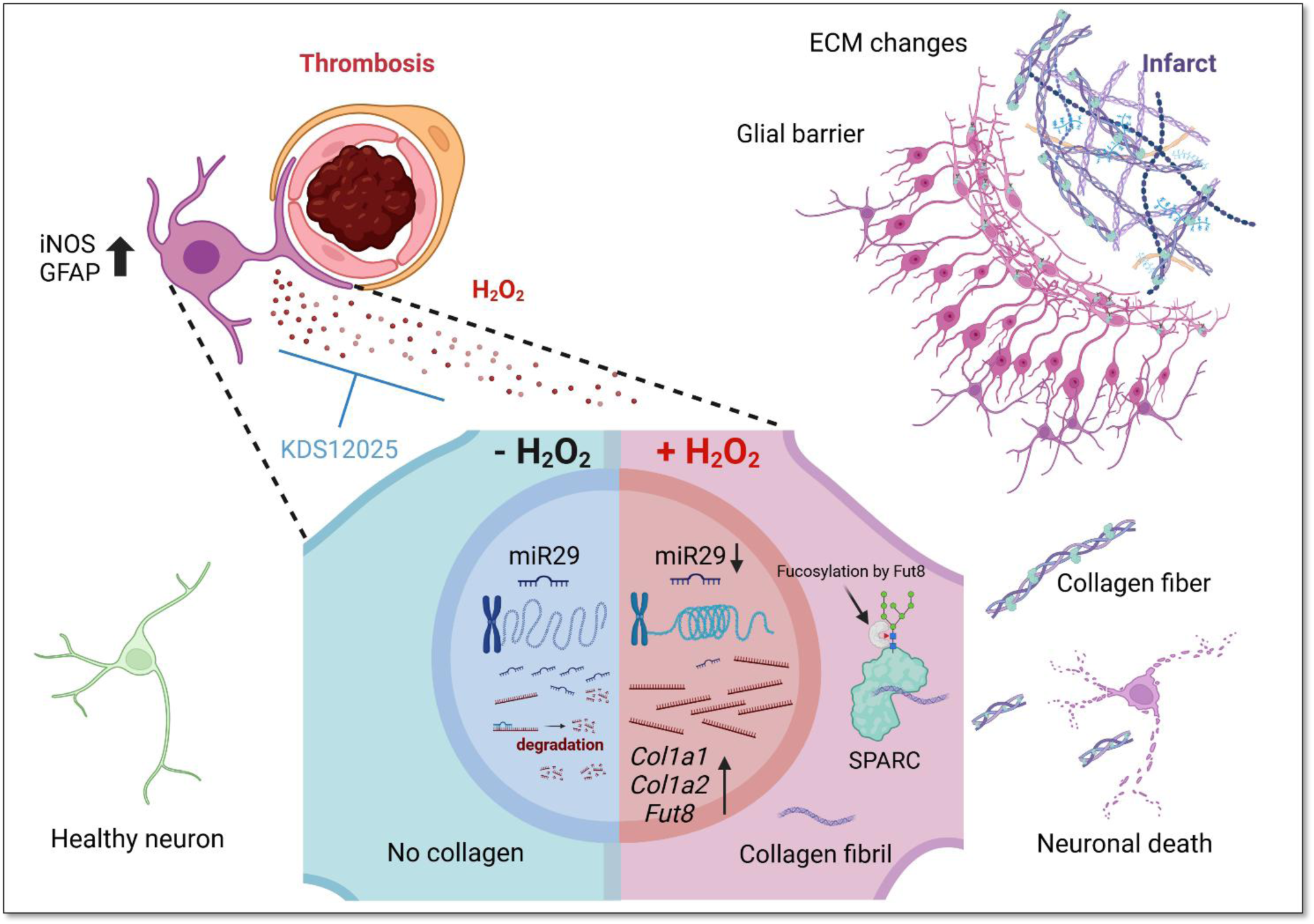
Molecular and cellular mechanism of astrocytic collagen production and neuronal death in PT-induced ischemic stroke. PT triggers excessive H_2_O_2_ production, leading to the upregulation of COL1-related genes in astrocytes. The chromatin region for miR-29, which normally degrades *Col1α1* mRNA, becomes less accessible. Consequently, the upregulated *Col1α1* and *Col1α2* form a collagen fibril triple helix that binds to N-glycosylated SPARC. This complex is then secreted, causing neuronal death and accumulating as collagen fibers. This accumulation may also trigger fibroblast-mediated fibrosis in the core region, leading to a fibrotic scar with a glial barrier.

**Table S1:**
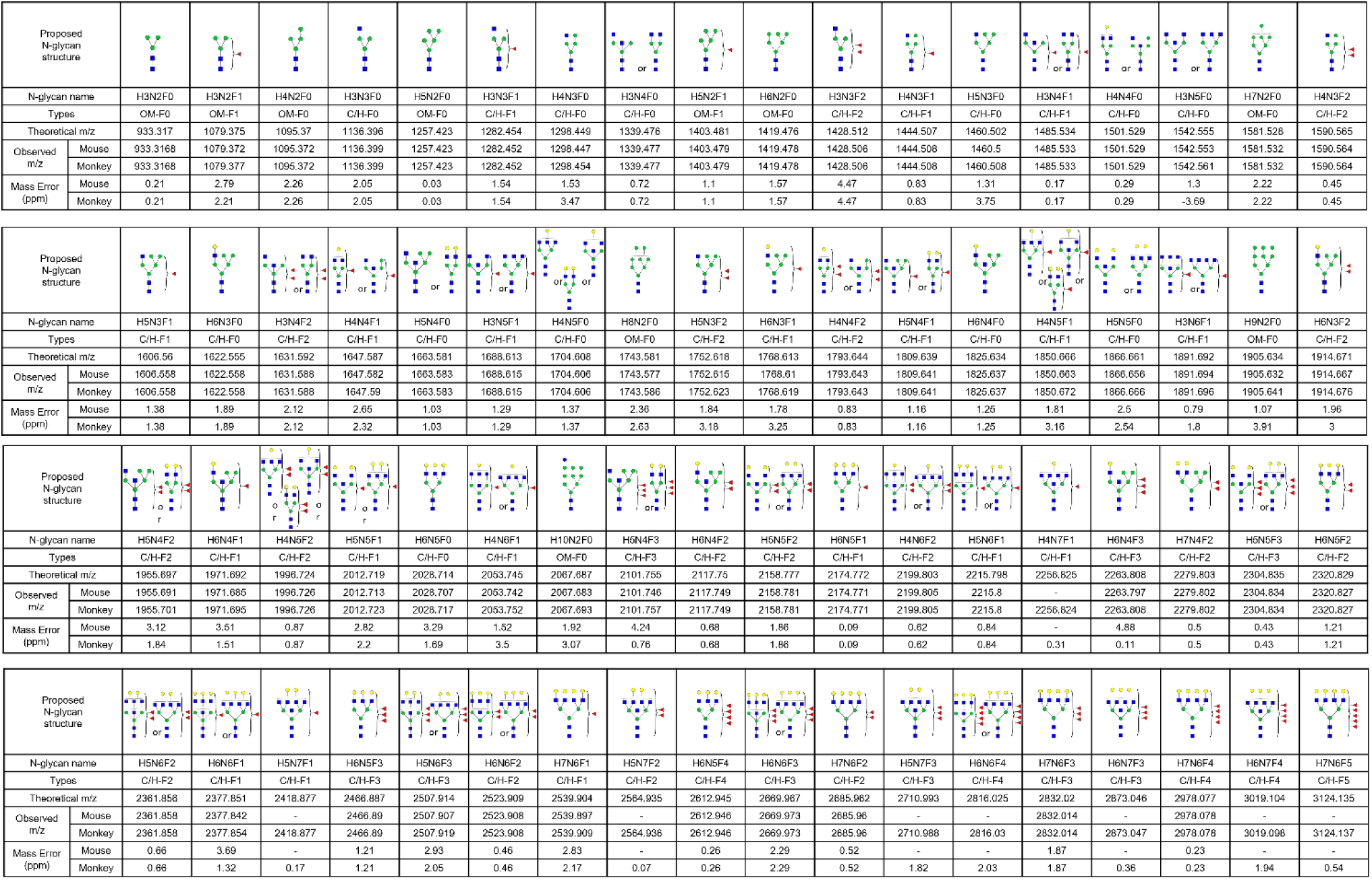
List of N-liked glycans measured in mouse and monkey brain. The proposed N-glycan structure represent by symbols, hexose (H), mannose (green circle), galactose (yellow circle), N-acetylglucosamine (GlcNAc, N, blue square), and fucose (F, red triangle) (left bottom). The measured N-glycans were classified into oligomannose (OM), complex or hybrid (C/H) types with fucosylated glycans. (F0: zero, F1: mono, F2: di, F3: tri, F4: tetra, F5: penta)

## Methods

### Study Approval

All animal care and experimental procedures were approved by the Institutional Animal Care and Use Committees (IACUC). For mice, the procedures were approved by the Institute for Basic Science (IBS; Daejeon, Republic of Korea, IBS-2023-010). For NHPs, the procedures were approved by the Daegu–Gyeongbuk Medical Innovation Foundation (KMEDI hub; Daegu, Republic of Korea, KMEDI-23080101-00).

### Animals

C57BL/6 mice were used for drug treatment experiments. GFAPcreERT2, Aldh1/1creERT2, Adlh1/1eGFP and Ai14 mice were used for astrocyte intra-vital imaging and astrocyte-specific gene silencing. All mice were weaned at P21 and housed in a specific pathogen-free controlled animal facility under a 12 h-12 h light-dark cycle of 21C and a 40-60% humidity range. Mice were allowed free access to food and water.

Two male and one female cynomolgus macaques (Macaca fascicularis), aged 3-4 years and weighing 2.6-5.1 kg, were used in the study. Animals were housed in pairs under controlled conditions (temperature: 25 ± 1°C, humidity: 50 ± 10%, 12-hour light/dark cycle) with ad libitum access to food and water.

### Cranial window surgery (CWS)

CWS was performed by our previous study^27^. Briefly, mice were deeply anesthetized with vaporized isoflurane and placed into stereotaxic frames (RWD). The scalp was incised, and a round hole (2-2.5 mm diameter) was made at the border of bregma including the M1, M2 cortical region using a hand drill. The hole was covered by a round glass coverslip (small diameter round cover glass, #1217N66, Harvard Apparatus) and was fixed by dental acrylic resin (1234, Lang dental). After implantation, mice were housed for 2-3 weeks to allow recovery from the surgery.

### Intra-vital imaging and ischemic stroke modeling by PT

Mice were anesthetized by intraperitoneal injection of Ketamine and Xylazine (10mg/kg) mixture and mounted on the stereotaxic module with a heating pad. 50 ul of Rose Bengal (15 mg/ml) was intravenously injected and 561 nm laser was irradiated for 100 seconds to induce cerebral infarction using an Intravital confocal microscope (IVIM-C, Korea). For tracking glial barrier formation, GFAPCreERT2 crossed with Ai14 mouse and Aldh1/1eGFP were imaged every week until a month at the same location.

NHPs were anesthetized with intramuscular injections of ketamine (8 mg/kg) and medetomidine (0.05 mg/kg), followed by maintenance with 1.2-1.5% isoflurane. A craniotomy was performed to expose the motor and premotor cortices. Rose Bengal (20 mg/mL) was administered intravenously at 80 mg/kg while a 561 nm laser was applied for 20 minutes to induce thrombosis. Post-surgery, animals received enrofloxacin (5 mg/kg), meloxicam (0.2 mg/kg), and tramadol (4 mg/kg) for postoperative management.

### Small molecule treatment

KDS12025, an H_2_O_2_ decomposing enhancer, was dissolved in 100 uL of DMSO and diluted in various concentrations by adding DPBS (10, 1, 0.1, 0.01, and 0.001 mg/kg). The final concentration was calculated according to individual body weight by adding normal saline. The drug was applied 10 minutes or 24 hours after thrombosis for three consecutive days by intraperitoneal (IP) administration. In the oral administration group, water consumption in each cage was calculated, and the proper concentration of KDS12025 was dissolved and applied to the water. In total, 36 mice were treated and were involved in experiments.

NHPs received KDS12025 by endotracheal intubation at 0.1 mg/kg for one day or 0.03 mg/kg for three days from 10 minutes post-PT by endotracheal intubation. After treatment, NHPs were moved to home cages and were monitored for adverse drug reactions for several hours by a veterinarian.

### Cylinder test

The cylinder test was employed to evaluate forelimb motor function. During the test, animals were placed in a transparent acrylic cylinder (10 cm in diameter and 20 cm in height) for 5 minutes to observe the frequency of forelimb usage. The number of ipsilateral and contralateral forepaw usages was recorded for each session. The test score was determined as the percentage of ipsilateral usage out of the total forepaw usages. The impaired forelimb usage was calculated using the following formula below.

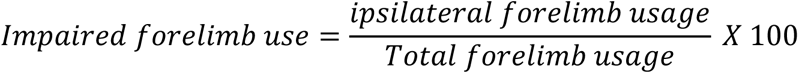

### Magnetic Resonance Imaging (MRI) acquisition and analysis

MRI data were acquired using a Siemens Skyra 3T scanner (Erlangen, Germany) with a gradient coil insert (XQ gradients; 45 mT/m maximum gradient strength, 200 mT/m/s maximum slew rate) and a 15-channel TX/RX coil. The lesion caused by photothrombosis were quantified with 3D Slicer (version 5.6.1). Tractography was performed with Generalized Q-sampling Imaging (GQI), and the corticospinal tract (CST) fiber count was calculated using DSI Studio (version Chen Jan 2024).

### Single Fruit Reaching Test (SFRT)

To evaluate hand function and fine motor skills in NHPs following PT, a single fruit reaching test (SFRT) was conducted. Animals underwent quarantine and acclimatization for the experimental chair, followed by baseline hand dominance assessment. Multiple reach-and-retrieve tasks were performed and recorded during each session, both before and after PT. Success was defined as retrieving and consuming the food within 10 seconds. Additionally, the type of grasp used—pinch grip (precise finger movements) versus power grip (using the entire hand)—was evaluated to assess the impact of the PT and the effect of KDS12025 on fine motor skills.

### Tissue Processing and Histological Assessment

The mice were anesthetized with isoflurane, and their brains were harvested following heart perfusion with normal saline (NS) and 4% paraformaldehyde (PFA) sequentially. The brains were then post-fixed in 4% PFA at 4°C overnight, followed by immersion in a 30% sucrose solution for at least 48 hours. Subsequently, coronal sections were cut at a thickness of 30µm.

For preparation of NHP, the animals were deeply anesthetized with a lethal dose of 5% Isoflurane. After the removal of blood by transcardial perfusion with 0.9% normal saline, the brain was harvested and fixed in 4% PFA in phosphate-buffered saline (PBS) and post-fixed in the same fixative overnight at 4°C to ensure thorough preservation of tissue architecture. The fixed brains were serially sectioned with brain matrices (15043, Ted Pella) and were prepared for paraffin (4 um) or cryosection (40 um).

### Immunofluorescence staining

Both mouse and NHP brain sections were blocked in 0.1 M PBS containing 0.3% Triton X-100 (Sigma) and 5% normal donkey serum (Genetex) for 1 hour at room temperature. The primary antibodies including GFAP (AB5541, 500:1), NeuN (3995040, 1:500), Collagen I (ab21286, 1:200), SPARC (33-5500, 1:200), Iba1 (019-19741, 1:500), and iNOS (ab15323, 1:500) in blocking solution were immunostained. The samples were incubated with the primary antibodies overnight at 4°C. After three washes in 0.1 M PBS, the sections were incubated with appropriate secondary antibodies for 1 hour. Following three additional washes in 0.1 M PBS and a 10-minute staining with DAPI at 1:1000 (Pierce), the samples were mounted on slides using a mounting solution (Vector, [catalog number]).

The sectioned brain of NHP was blocked with the solution containing 5% normal donkey serum with 0.3% Triton-X in 1X PBS for 1 hour, then incubated in a primary antibody cocktail including GFAP (AB5541, 500:1), NeuN (3995040, 1:500), and Collagen I (ab21286, 1:200) at 4C rocker overnight. The next day, the sample was washed 3 times in 1X PBS, and relevant secondary antibodies were applied for 1 h at room temperature. After 2 times washing, DAPI (1:1000, Dako, 62248) was stained, and samples were mounted on a slide with the mounting solution (Dako, F4680).

Images were captured using a Zeiss LSM 900 confocal microscope. For morphological analysis, 30-40 stacks with a 1 um interval were acquired.

### Astrocyte surface and Sholl analysis with Imaris

Astrocyte surface and Sholl analysis in both mouse and NHP were conducted following the methodology detailed in our previous report^11^. In brief, the astrocyte filaments and dendrites from the maximally projected confocal image were reconstructed using Imaris software. Specific parameters were adjusted to visualize the GFAP signal from the channel, allowing for the extraction of volume, length, branch depth, and branch points. Sholl analysis was carried out using the ImageJ plug-in, which automatically drew a series of concentric circles at 10 µm intervals from the center of the DAPI signal.

### Cortical astrocyte culture and H_2_O_2_ treatment

Primary cortical astrocytes were harvested as described ^44^. Briefly, cerebral cortices of P4 mice were dissected free of meninges, and cortices were mechanically dissociated with pipet. Cells were plated on 6 or 24 well plates, coated with 0.1 mg/ml of poly-D-lysine (Sigma). Cultured astrocytes were maintained in Dulbecco’s modified Eagle’s medium (DMEM, Corning) supplemented with 4.5 g/L glucose, L-glutamine, sodium pyruvate, 10% heat-inactivated horse serum, 10% heat-inactivated fetal bovine serum (FBS), and 1000 units/ml of penicillin-streptomycin at 37℃ with 5% CO2. Three days after culture, cells were washed with pipetting, and media was replaced to eliminate dead cells and debris. 200 µm of H_2_O_2_ (Sigma, 88597) was treated for 3 hours and washed with maintaining media. After 24 or 72 hours of washing, cells were harvested for various analyses, including immunocytochemistry, western blotting, qRT-PCR, and RNA and ATAC-seq.

### Sandwich culture

For the sandwich culture experiment, we used a 6-well culture plate with inserts (SPLinsert, #35006, SPL Life Sciences) designed to co-culture cortical neurons and astrocytes without direct contact. Primary cortical astrocytes and neurons were seeded separately in the plate and insert, respectively. Astrocytes were treated with 200 µM H₂O₂ for 3 hours, followed by washing with maintenance media. After 24 hours, the inserts containing astrocytes were placed onto the 6-well plate containing the seeded neurons, completing the sandwich culture setup.

### Preparation of gene-specific shRNA and shRNA virus

The shRNA sequences for Scrambled, *Col1α1*, and *Fut8* were designed using BLOCK-iT RNAi Designer (Invitrogen) and then cloned into pSico and pSicoR Adeno-Associated Virus (AAV) vectors with PhP.eB serotype as previously described^29^. pSico and pSicoR vectors were chosen for Cre-specific or plasmid-based shRNA expression both in vitro and in vivo. Each construct was sequenced to verify the cloning accuracy. The cloned shRNA constructs were packaged into AAV viruses at the IBS Virus Facility (http://ibs.re.kr/virusfacility). The effectiveness of each shRNA was assessed in cortical astrocytes by measuring mRNA levels with qRT-PCR. The sequences for each target are listed in supplementary figures.

### RT-qPCR

RT-qPCR was conducted with SYBR Green PCR Master Mix as previously described^14^. In short, reactions were set up in triplicate with a total volume of 10 μl, including 10 pM primer, 4 μl cDNA, and 5 μl Power SYBR Green PCR Master Mix (Applied Biosystems). The mRNA levels of each gene were normalized to 18S mRNA, and fold changes were calculated using the 2−ΔΔCT method. The primer sequences used for RT-qPCR were as follows:

> *Col1α1* forward 5′-GCC AAGAAGACATCCCTGAAG-3′,

> *Col1α1* Reverse 5′-TGT GGCAGATACAGATCAAGC-3′

> Gapdh forward 5′-CCATGG AGAAGGCTGGGG-3′,

> Gapdh Reverse 5′-CAACAG CGGTGCAGGTAAGTG-3′

> *Fut8* forward 5′-GCTTGAACGCTTAAAACAGCA-3′,

> *Fut8* Reverse 5′-AATGGGGCCTTCTGGTATTC-3′

### Cortical neuron culture, sandwich-culture, and viability assay

Cerebral cortices from E18 mouse pups were dissected and dissociated into single cells in Hank’s balanced salt solution (HBSS, Welgene). The cells were then centrifuged for 2 minutes at 2000 rpm, and the resulting cell pellet was resuspended in Dulbecco’s modified Eagle’s medium (DMEM, Welgene) containing 10% fetal bovine serum (FBS, Welgene) and 1% penicillin-streptomycin (Gibco). The cells were plated on 10 mm coverslips coated with poly-D-lysine (0.1 mg/mL in deionized water, Sigma). They were placed in a 24-well plate and incubated at 37°C in a humidified atmosphere with 5% CO2, with the medium being replaced every 3-4 days. After 10 days, the cells were treated with various doses of Col1 or sandwich-cultured with astrocytes treated with H2O2. Forty-eight hours after co-culture, the neurons were incubated in phosphate-buffered saline (PBS) containing 2 µM calcein AM for 15 minutes at room temperature. After staining, the samples were washed twice with PBS and imaged using a fluorescence microscope.

### Immunocytochemistry

Cells were fixed with 4% paraformaldehyde (PFA) for 15 minutes and merged in a blocking solution containing 0.3% triton X-100 with 5% normal donkey serum in PBS for 15 minutes at room temperature. Then, cells were incubated overnight with primary antibodies, including GFAP (AB5541, 500:1) and Collagen I (ab21286, 1:200) at 4C, followed by 3 times of washes and incubation with a specific secondary antibody for 30 minutes at room temperature. After 2 times of washing, cell nuclei were stained with DAPI. Image was taken using ZIESS LSM-900 microscope.

### Western blot analysis

Cells were harvested with ice-cold RIPA buffer containing 1X of PPI (Sigma, PPC-1010) and incubated for 1 hour at 4C. lysates were centrifuged with 15,000 RPM for 10 minutes in 4C and the supernatant was harvested. Collected proteins were denatured and loaded on pre-made SDS-gel (Bio-rad, 4566033).

Transferred blots were incubated in a blocking solution containing 4% skim milk and washed with 1X TBS-T 3 times. Then, the blots were incubated with the following primary antibodies: Collagen I (ab21286, 1:500) and GAPDH (ab8245, 5000:1) at 4C for 24 h on the rocker. After 3 times of washing with TBS-T, the blots were incubated with the appropriate secondary antibodies conjugated to horseradish peroxidase at room temperature for 2h. Then, the blots were developed by Immobilon Western ECL solution (WBKLS0500, Merck Millipore), and immunoreactive bands were imaged using an Image Station 4000MM (745280, Kodak).

### Illumina library preparation and sequencing

Sample libraries were prepared by NEBNext Ultra^TM^ Ⅱ RNA Library prep kit for Illumina (E7770, NEB), Multiplex Oligos, and Dynabeads mRNA DIRECT Kit (61012, Invitrogen) according to the manufacturer’s instructions.

### Bulk RNA sequencing data processing and analysis

Quality evaluation is performed using FASTQC (v.0.12.1), and adaptor trimming is performed using Cutadapt (v.2.6) with the following options (-q 10 -m 15 -e 0.10). Trimmed reads are mapped to the mouse genome (GRCm38/mm10) using STAR (v.2.7.3a), and features are counted with Subread. Differential expression analysis is conducted using DESeq2. Genes are considered significantly differentially expressed if the P-value is < 0.05. PCA is performed on VST-transformed values (implemented in DESeq2). Heatmaps are generated from z-score transformed normalized counts. To identify pathways and functions associated with different cellular treatment conditions, differentially expressed genes (DEGs) showing significant changes are selected based on DEG analysis (calculated using log2 fold change and P-values). GO and pathway enrichment analyses are then performed using enrichR.

### ATAC-Seq data preprocessing and analysis

Paired-end ATAC-seq reads are first assessed using FASTQC (v.0.12.1). Reads are trimmed using Cutadapt (v.2.6) with the options (-q 10 -m 15 -e 0.10). Paired-end reads are aligned against the GRCm38/mm10 mouse genome assembly with Bowtie2 (v2.2.5) in local mode, sensitive settings, and a maximum fragment size of 2,000. Duplicated reads are marked using Biobambam (v.2.0.87). Alignments are filtered with SAMtools (v.1.6) to exclude reads with a mapping quality of <30, duplicates, and those not properly paired. ATAC-seq peaks are called for each replicate using MACS2 with the BAMPE format. Parameters include shifting reads by -100 bp (--shift -100) and extending reads by 200 bp (--extsize 200). Consensus peaks are generated by merging data (.narrowPeak) and consolidating overlaps using the GenomicRanges package in R. For downstream analysis, annotations and distribution analyses of consensus peaks are performed using the ChIPseeker package in R. Motif analysis is conducted using the FIMO tool from the MEME suite.

### Matrix-assisted laser desorption/ionization mass spectrometry Imaging (MALDI-MSI) for spatial distribution measurement of N-glycans

For the sample preparation method for MALDI-MSI of N-glycans, refer to the previously published paper^58^. Mouse brains fixed with 4% paraformaldehyde (PFA) were cut into coronal sections at a thickness of 14 μm using a CM1950 cryostat (Leica) and then thaw-mounted on conductive indium tin oxide (ITO) slides (Applied Surface Tech Asia). The NHP brains, fixed with 4% PFA, were embedded in paraffin and attached to ITO slides at a thickness of 5 μm. 0.1 mg/ml PNGase F PRIME-LY™ ULTRA GLYCOSIDASE (N-zyme Scientifics) and 7 mg/ml α-cyano-4-hydroxycinnamic acid (CHCA; Sigma Aldrich) were applied to the sample slide using an M3+ or M5 sprayer. N-glycan images of the mouse brain were obtained using the MALDI QTOF instrument, timsTOF fleX (Bruker Daltonics). Mass spectra were recorded using 500 laser shots per pixel in positive ion mode. Laser focusing size and spatial resolution were set to 20 μm for mice and 50 μm for NHP brain tissues. The measured mass spectra were normalized to total ion count (TIC) and converted to mass spectrometry images in SCiLS Lab 2023a software (Bruker Daltonics). The extracted data from SCiLS software were analyzed by one-way ANOVA in GraphPad Prism (version. 10.2.0) and uploaded to MetaboAnalyst (version. 6.0) for partial least squares-discriminant analysis (PLS-DA), variable importance in projection (VIP) analysis, and hierarchical clustering heatmaps. The structure of the proposed n-glycan was annotated through GlycoWorkBench based on previously published papers^59^. The extended table contains a list of all N-glycans reported in this study and proposed N-glycan structures^60^ (Table S1).

### High-resolution serial optical coherence microscopy (SOCM) imaging and 3D reconstruction

Serial Optical Coherence Microscopy (SOCM) is an imaging technique that integrates spectral-domain SOCM with vibratome-based tissue sectioning, facilitating three-dimensional quantification of tissues without requiring labeling or staining ^34^. We performed Serial SOCM imaging for fibrotic scar quantification. A mosaic-based OCM approach was employed, alternating between tissue sectioning and imaging to acquire a large field-of-view image set, which was subsequently stacked to create a volumetric image of the mouse brain by sectioning with 30 µm of thickness^34,61^. The stage movement and camera acquisition processes are synchronized, enabling the region of interest to be automatically acquired in a mosaic manner using LabVIEW software. The 3D image reconstruction utilized custom-developed registration software, and scar regions were segmented and labeled using Amira software. Finally, segmented scars were quantified by calculating voxel values in the 3D images to determine their volume measurements.

### Preparation of brain slice

Mice were anesthetized with isoflurane and subsequently decapitated for brain extraction. Brains were rapidly placed in an ice-cold slicing solution containing 234 mM sucrose, 2.5 mM KCl, 104 mM MgSO₄, 1.25 mM NaH₂PO₄, 24 mM NaHCO₃, 0.5 mM CaCl₂·2H₂O, and 11 mM glucose. Coronal brain slices (300 µm thick) were prepared using a vibrating microtome (LinearSlicer PRO7N, DSK, Japan) and transferred to artificial cerebrospinal fluid (aCSF) containing 126 mM NaCl, 24 mM NaHCO₃, 1 mM NaH₂PO₄, 2.5 mM KCl, 2.5 mM CaCl₂, 2 mM MgCl₂, and 10 mM D-glucose (pH 7.4). Slices were incubated at room temperature for a minimum of 30 minutes before electrophysiological recordings. For recording, slices were placed in a chamber perfused continuously with aCSF at a rate of 2–3 mL/min using a peristaltic pump (Miniplus 3, Gilson). The chamber was mounted on an upright microscope (Nikon) equipped with a 40× water-immersion objective (NA 1.00, WD 2.80 mm) and infrared differential interference contrast (IR-DIC) optics. Neuronal morphology was visualized with an sCMOS camera (Hamamatsu) and Imaging Workbench software (INDEC BioSystems). Whole-cell voltage-clamp recordings were obtained from neurons within the infarcted region, with a holding potential set to –70 mV.

### Neuron characterization and spontaneous EPSCs and IPSCs measurement

Whole-cell patch-clamp recordings were performed using glass pipette electrodes with a resistance of 5–7 MΩ. For neuronal characterization, pipettes were filled with an internal solution containing 120 mM potassium gluconate, 10 mM KCl, 1 mM MgCl₂, 0.5 mM EGTA, and 30 mM HEPES (pH adjusted to 7.2). Electrical signals were digitized and sampled at 20 kHz (every 50 µs) using a Digidata 1550B digitizer and a Multiclamp 700B amplifier (Molecular Devices), controlled by pCLAMP 11.2 software. Signals were low-pass filtered at 2 kHz.

Spontaneous excitatory and inhibitory postsynaptic currents (sEPSCs and sIPSCs) were recorded for 4–5 minutes. For these recordings, the internal pipette solution contained 140 mM Cs-methanesulfonate, 8 mM NaCl, 1 mM MgCl₂, 0.5 mM EGTA, 10 mM HEPES, 7 mM phosphocreatine di(tris) salt, 4 mM Mg-ATP, 0.3 mM Na₂-GTP, and 5 mM QX-314, with the pH adjusted to 7.3 using NMDG. To record sEPSCs, cells were voltage-clamped at –70 mV to enhance detection of inward glutamatergic currents while minimizing inhibitory currents. For sIPSC recordings, the same internal solution was used, and cells were held at 0 mV to isolate GABA_A receptor-mediated Cl⁻ currents. Recorded traces were analyzed using MiniAnalysis software (version 7; Synaptosoft, BlueCell, Seoul, Korea).

### Statistics and reproducibility

In this paper, data were presented as the mean ± s.e.m. Statistical analyses were carried out using Prism v.10 (GraphPad Software). To compare the two groups, either an unpaired t-test or a Mann-Whitney U test was used, based on whether the data met the normality assumption. For comparisons between groups with more than two time points, one-way or two-way ANOVA followed by Dunnett’s post hoc test was used, as detailed in the figure legends. Statistical significance was defined as p < 0.05 and detailed in the figure legends. Further statistical information is available in Supplementary Table 2 and has been deposited in a public repository.

## References

1. Massaad, C.A., and Klann, E. (2011). Reactive oxygen species in the regulation of synaptic plasticity and memory. Antioxid Redox Signal 14, 2013–2054. 10.1089/ars.2010.3208.

2. Hu, Q., Khanna, P., Ee Wong, B.S., Lin Heng, Z.S., Subhramanyam, C.S., Thanga, L.Z., Sing Tan, S.W., and Baeg, G.H. (2018). Oxidative stress promotes exit from the stem cell state and spontaneous neuronal differentiation. Oncotarget 9, 4223–4238. 10.18632/oncotarget.23786.

3. Yang, Y., Bazhin, A.V., Werner, J., and Karakhanova, S. (2013). Reactive oxygen species in the immune system. Int Rev Immunol 32, 249–270. 10.3109/08830185.2012.755176.

4. Kakizawa, S., Arasaki, T., Yoshida, A., Sato, A., Takino, Y., Ishigami, A., Akaike, T., Yanai, S., and Endo, S. (2024). Essential role of ROS – 8-Nitro-cGMP signaling in long-term memory of motor learning and cerebellar synaptic plasticity. Redox Biology 70, 103053. 10.1016/j.redox.2024.103053.

5. Su, L.J., Zhang, J.H., Gomez, H., Murugan, R., Hong, X., Xu, D., Jiang, F., and Peng, Z.Y. (2019). Reactive Oxygen Species-Induced Lipid Peroxidation in Apoptosis, Autophagy, and Ferroptosis. Oxid Med Cell Longev 2019, 5080843 10.1155/2019/5080843.

6. Sultan, S., Li, L., Moss, J., Petrelli, F., Cassé, F., Gebara, E., Lopatar, J., Pfrieger, F.W., Bezzi, P., and Bischofberger, J. (2015). Synaptic integration of adult-born hippocampal neurons is locally controlled by astrocytes. Neuron 88, 957–972.

7. Shadfar, S., Parakh, S., Jamali, M.S., and Atkin, J.D. (2023). Redox dysregulation as a driver for DNA damage and its relationship to neurodegenerative diseases. Translational Neurodegeneration 12, 18 10.1186/s40035-023-00350-4.

8. Wang, X., and Michaelis, E.K. (2010). Selective neuronal vulnerability to oxidative stress in the brain. Frontiers in aging neuroscience 2, 1224.

9. Franzoni, F., Scarfò, G., Guidotti, S., Fusi, J., Asomov, M., and Pruneti, C. (2021). Oxidative stress and cognitive decline: the neuroprotective role of natural antioxidants. Frontiers in neuroscience 15, 729757.

10. Rodrigo, R., Fernández-Gajardo, R., Gutiérrez, R., Matamala, J.M., Carrasco, R., Miranda-Merchak, A., and Feuerhake, W. (2013). Oxidative stress and pathophysiology of ischemic stroke: novel therapeutic opportunities. CNS Neurol Disord Drug Targets 12, 698–714. 10.2174/1871527311312050015.

11. Chun, H., Im, H., Kang, Y.J., Kim, Y., Shin, J.H., Won, W., Lim, J., Ju, Y., Park, Y.M., Kim, S., et al. (2020). Severe reactive astrocytes precipitate pathological hallmarks of Alzheimer’s disease via H(2)O(2)(-) production. Nat Neurosci 23, 1555–1566. 10.1038/s41593-020-00735-y.

12. Dias, V., Junn, E., and Mouradian, M.M. (2013). The role of oxidative stress in Parkinson’s disease. J Parkinsons Dis 3, 461–491. 10.3233/jpd-130230.

13. Escartin, C., Galea, E., Lakatos, A., O’Callaghan, J.P., Petzold, G.C., Serrano-Pozo, A., Steinhäuser, C., Volterra, A., Carmignoto, G., and Agarwal, A. (2021). Reactive astrocyte nomenclature, definitions, and future directions. Nature neuroscience 24, 312–325.

14. Sofroniew, M.V. (2020). Astrocyte reactivity: subtypes, states, and functions in CNS innate immunity. Trends in immunology 41, 758–770.

15. Sofroniew, M.V. (2009). Molecular dissection of reactive astrogliosis and glial scar formation. Trends Neurosci 32, 638–647. 10.1016/j.tins.2009.08.002.

16. Haist, V., Ulrich, R., Kalkuhl, A., Deschl, U., and Baumgärtner, W. (2012). Distinct spatio-temporal extracellular matrix accumulation within demyelinated spinal cord lesions in Theiler’s murine encephalomyelitis. Brain Pathol 22, 188–204. 10.1111/j.1750-3639.2011.00518.x.

17. Matafora, V., Gorb, A., Yang, F., Noble, W., Bachi, A., Perez-Nievas, B.G., and Jimenez-Sanchez, M. (2023). Proteomics of the astrocyte secretome reveals changes in their response to soluble oligomeric Aβ. J Neurochem 166, 346–366. 10.1111/jnc.15875.

18. Jucker, M., Bialobok, P., Kleinman, H.K., Walker, L.C., Hagg, T., and Ingram, D.K. (1993). Laminin-like and laminin-binding protein-like immunoreactive astrocytes in rat hippocampus after transient ischemia. Antibody to laminin-binding protein is a sensitive marker of neural injury and degeneration. Ann N Y Acad Sci 679, 245–252. 10.1111/j.1749-6632.1993.tb18304.x.

19. Ji, K., and Tsirka, S.E. (2012). Inflammation modulates expression of laminin in the central nervous system following ischemic injury. J Neuroinflammation 9, 159. 10.1186/1742-2094-9-159.

20. Verkhratsky, A., Butt, A., Li, B., Illes, P., Zorec, R., Semyanov, A., Tang, Y., and Sofroniew, M.V. (2023). Astrocytes in human central nervous system diseases: a frontier for new therapies. Signal Transduct Target Ther 8, 396. 10.1038/s41392-023-01628-9.

21. Won, W., Bhalla, M., Lee, J.H., and Lee, C.J. (2025). Astrocytes as Key Regulators of Neural Signaling in Health and Disease. Annu Rev Neurosci. 10.1146/annurev-neuro-112723-035356.

22. Macfarlane, L.A., and Murphy, P.R. (2010). MicroRNA: Biogenesis, Function and Role in Cancer. Curr Genomics 11, 537–561. 10.2174/138920210793175895.

23. Maurer, B., Stanczyk, J., Jüngel, A., Akhmetshina, A., Trenkmann, M., Brock, M., Kowal-Bielecka, O., Gay, R.E., Michel, B.A., Distler, J.H., et al. (2010). MicroRNA-29, a key regulator of collagen expression in systemic sclerosis. Arthritis Rheum 62, 1733–1743. 10.1002/art.27443.

24. Agarwal, V., Bell, G.W., Nam, J.W., and Bartel, D.P. (2015). Predicting effective microRNA target sites in mammalian mRNAs. Elife 4. 10.7554/eLife.05005.

25. Boraschi-Diaz, I., Wang, J., Mort, J.S., and Komarova, S.V. (2017). Collagen type I as a ligand for receptor-mediated signaling. Frontiers in Physics 5, 12.

26. Dias, D.O., and Göritz, C. (2018). Fibrotic scarring following lesions to the central nervous system. Matrix Biol 68-69, 561-570. 10.1016/j.matbio.2018.02.009.

27. Lee, J., Kim, J.G., Hong, S., Kim, Y.S., Ahn, S., Kim, R., Chun, H., Park, K.D., Jeong, Y., Kim, D.E., et al. (2022). Longitudinal intravital imaging of cerebral microinfarction reveals a dynamic astrocyte reaction leading to glial scar formation. Glia 70, 975–988. 10.1002/glia.24151.

28. Berndt, A., Lee, J., Won, W., Kimball, K., Neiswanger, C., Schattauer, S., Wang, Y., Yeboah, F., Ruiz, M., Evitts, K., et al. (2024). Ultra-fast genetically encoded sensor for precise real-time monitoring of physiological and pathophysiological peroxide dynamics. Res Sq. 10.21203/rs.3.rs-4048855/v1.

29. Chan, K.Y., Jang, M.J., Yoo, B.B., Greenbaum, A., Ravi, N., Wu, W.L., Sánchez-Guardado, L., Lois, C., Mazmanian, S.K., Deverman, B.E., and Gradinaru, V. (2017). Engineered AAVs for efficient noninvasive gene delivery to the central and peripheral nervous systems. Nat Neurosci 20, 1172–1179. 10.1038/nn.4593.

30. dos Santos, N.V., Saponi, C.F., Ryan, T.M., Primo, F.L., Greaves, T.L., and Pereira, J.F.B. (2020). Reversible and irreversible fluorescence activity of the Enhanced Green Fluorescent Protein in pH: Insights for the development of pH-biosensors. International Journal of Biological Macromolecules 164, 3474–3484. 10.1016/j.ijbiomac.2020.08.224.

31. Sheng, W.S., Hu, S., Feng, A., and Rock, R.B. (2013). Reactive oxygen species from human astrocytes induced functional impairment and oxidative damage. Neurochem Res 38, 2148–2159. 10.1007/s11064-013-1123-z.

32. Trombetta-Esilva, J., and Bradshaw, A.D. (2012). The Function of SPARC as a Mediator of Fibrosis. Open Rheumatol J 6, 146–155. 10.2174/1874312901206010146.

33. Bradshaw, A.D. (2009). The role of SPARC in extracellular matrix assembly. J Cell Commun Signal 3, 239–246. 10.1007/s12079-009-0062-6.

34. Min, E., Ban, S., Lee, J., Vavilin, A., Baek, S., Jung, S., Ahn, Y., Park, K., Shin, S., Han, S., et al. (2020). Serial optical coherence microscopy for label-free volumetric histopathology. Scientific Reports 10, 6711. 10.1038/s41598-020-63460-3.

35. Kawamata, T., Dietrich, W.D., Schallert, T., Gotts, J.E., Cocke, R.R., Benowitz, L.I., and Finklestein, S.P. (1997). Intracisternal basic fibroblast growth factor enhances functional recovery and up-regulates the expression of a molecular marker of neuronal sprouting following focal cerebral infarction. Proc Natl Acad Sci U S A 94, 8179–8184. 10.1073/pnas.94.15.8179.

36. Brenneman, M., Sharma, S., Harting, M., Strong, R., Cox, C.S., Jr., Aronowski, J., Grotta, J.C., and Savitz, S.I. (2010). Autologous bone marrow mononuclear cells enhance recovery after acute ischemic stroke in young and middle-aged rats. J Cereb Blood Flow Metab 30, 140–149. 10.1038/jcbfm.2009.198.

37. Nam, M.H., Cho, J., Kwon, D.H., Park, J.Y., Woo, J., Lee, J.M., Lee, S., Ko, H.Y., Won, W., Kim, R.G., et al. (2020). Excessive Astrocytic GABA Causes Cortical Hypometabolism and Impedes Functional Recovery after Subcortical Stroke. Cell Rep 32, 107861. 10.1016/j.celrep.2020.107861.

38. Park, Y.M., Chun, H., Shin, J.I., and Lee, C.J. (2018). Astrocyte Specificity and Coverage of hGFAP-CreERT2 [Tg(GFAP-Cre/ERT2)13Kdmc] Mouse Line in Various Brain Regions. Exp Neurobiol 27, 508–525. 10.5607/en.2018.27.6.508.

39. Srinivasan, R., Lu, T.Y., Chai, H., Xu, J., Huang, B.S., Golshani, P., Coppola, G., and Khakh, B.S. (2016). New Transgenic Mouse Lines for Selectively Targeting Astrocytes and Studying Calcium Signals in Astrocyte Processes In Situ and In Vivo. Neuron 92, 1181–1195. 10.1016/j.neuron.2016.11.030.

40. Won, W., Lee, E.H., Gotina, L., Chun, H., Park, U., Kim, D., Kim, T.Y., Choi, J., Kim, Y., Park, S.J., et al. (2024). Astrocytic hemoglobin is an H_2_O_2_-decomposing peroxidase and therapeutic target for Alzheimer’s disease. bioRxiv, 2024.2005.2021.594979. 10.1101/2024.05.21.594979.

41. Porritt, M.J., Andersson, H.C., Hou, L., Nilsson, Å., Pekna, M., Pekny, M., and Nilsson, M. (2012). Photothrombosis-induced infarction of the mouse cerebral cortex is not affected by the Nrf2-activator sulforaphane. PloS one 7, e41090.

42. Wang, S.-H., Wu, T.-J., Lee, C.-W., and Yu, J. (2020). Dissecting the conformation of glycans and their interactions with proteins. Journal of biomedical science 27, 93.

43. Wu, T.J., Wang, S.H., Chen, E.S., Tsai, H.H., Chang, Y.C., Tseng, Y.H., and Yu, J. (2022). Loss of core-fucosylation of SPARC impairs collagen binding and contributes to COPD. Cell Mol Life Sci 79, 348. 10.1007/s00018-022-04381-4.

44. Murata, A., Gallese, V., Luppino, G., Kaseda, M., and Sakata, H. (2000). Selectivity for the shape, size, and orientation of objects for grasping in neurons of monkey parietal area AIP. Journal of neurophysiology 83, 2580–2601.

45. Castiello, U., and Begliomini, C. (2008). The cortical control of visually guided grasping. The Neuroscientist 14, 157–170.

46. Xu, Y., Gurusiddappa, S., Rich, R.L., Owens, R.T., Keene, D.R., Mayne, R., Höök, A., and Höök, M. (2000). Multiple binding sites in collagen type I for the integrins alpha1beta1 and alpha2beta1. J Biol Chem 275, 38981–38989. 10.1074/jbc.M007668200.

47. Wareham, L.K., Baratta, R.O., Del Buono, B.J., Schlumpf, E., and Calkins, D.J. (2024). Collagen in the central nervous system: contributions to neurodegeneration and promise as a therapeutic target. Mol Neurodegener 19, 11. 10.1186/s13024-024-00704-0.

48. Holl, D., Hau, W.F., Julien, A., Banitalebi, S., Kalkitsas, J., Savant, S., Llorens-Bobadilla, E., Herault, Y., Pavlovic, G., Amiry-Moghaddam, M., et al. (2024). Distinct origin and region-dependent contribution of stromal fibroblasts to fibrosis following traumatic injury in mice. Nature Neuroscience 27, 1285–1298. 10.1038/s41593-024-01678-4.

49. Franze, K., Janmey, P.A., and Guck, J. (2013). Mechanics in neuronal development and repair. Annual review of biomedical engineering 15, 227–251.

50. Silver, J., and Miller, J.H. (2004). Regeneration beyond the glial scar. Nature Reviews Neuroscience 5, 146–156. 10.1038/nrn1326.

51. Liu, D., Zhao, Z., Wang, A., Ge, S., Wang, H., Zhang, X., Sun, Q., Cao, W., Sun, M., Wu, L., et al. (2018). Ischemic stroke is associated with the pro-inflammatory potential of N-glycosylated immunoglobulin G. J Neuroinflammation 15, 123. 10.1186/s12974-018-1161-1.

52. Wen, B., Xu, K., Huang, R., Jiang, T., Wang, J., Chen, J., Chen, J., and He, B. (2022). Preserving mitochondrial function by inhibiting GRP75 ameliorates neuron injury under ischemic stroke. Mol Med Rep 25. 10.3892/mmr.2022.12681.

53. Anderson, M.A., Burda, J.E., Ren, Y., Ao, Y., O’Shea, T.M., Kawaguchi, R., Coppola, G., Khakh, B.S., Deming, T.J., and Sofroniew, M.V. (2016). Astrocyte scar formation aids central nervous system axon regeneration. Nature 532, 195–200. 10.1038/nature17623.

54. Olmez, I., and Ozyurt, H. (2012). Reactive oxygen species and ischemic cerebrovascular disease. Neurochem Int 60, 208–212. 10.1016/j.neuint.2011.11.009.

55. Yamato, M., Egashira, T., and Utsumi, H. (2003). Application of in vivo ESR spectroscopy to measurement of cerebrovascular ROS generation in stroke. Free Radical Biology and Medicine 35, 1619–1631.

56. Peters, O., Back, T., Lindauer, U., Busch, C., Megow, D., Dreier, J., and Dirnagl, U. (1998). Increased formation of reactive oxygen species after permanent and reversible middle cerebral artery occlusion in the rat. Journal of cerebral blood flow & metabolism 18, 196–205.

57. Forman, H.J., and Zhang, H. (2021). Targeting oxidative stress in disease: promise and limitations of antioxidant therapy. Nat Rev Drug Discov 20, 689–709. 10.1038/s41573-021-00233-1.

58. Drake, R.R., Powers, T.W., Norris-Caneda, K., Mehta, A.S., and Angel, P.M. (2018). In Situ Imaging of N-Glycans by MALDI Imaging Mass Spectrometry of Fresh or Formalin-Fixed Paraffin-Embedded Tissue. Curr Protoc Protein Sci 94, e68. 10.1002/cpps.68.

59. Ceroni, A., Maass, K., Geyer, H., Geyer, R., Dell, A., and Haslam, S.M. (2008). GlycoWorkbench: A Tool for the Computer-Assisted Annotation of Mass Spectra of Glycans. Journal of Proteome Research 7, 1650–1659. 10.1021/pr7008252.

60. Lee, J., Ha, S., Kim, M., Kim, S.-W., Yun, J., Ozcan, S., Hwang, H., Ji, I.J., Yin, D., and Webster, M.J. (2020). Spatial and temporal diversity of glycome expression in mammalian brain. Proceedings of the National Academy of Sciences 117, 28743–28753.

61. Min, E., Lee, J., Vavilin, A., Jung, S., Shin, S., Kim, J., and Jung, W. (2015). Wide-field optical coherence microscopy of the mouse brain slice. Opt. Lett. 40, 4420–4423. 10.1364/OL.40.004420.

